# Nanobodies combined with DNA-PAINT super-resolution reveal a staggered titin nano-architecture in flight muscles

**DOI:** 10.1101/2022.04.14.488306

**Authors:** Florian Schueder, Pierre Mangeol, Eunice HoYee Chan, Renate Rees, Jürgen Schünemann, Ralf Jungmann, Dirk Görlich, Frank Schnorrer

## Abstract

Sarcomeres are the force producing units of all striated muscles. Their nanoarchitecture critically depends on the large titin protein, which in vertebrates spans from the sarcomeric Z-disc to the M-band and hence links actin and myosin filaments stably together. This ensures sarcomeric integrity and determines the length of vertebrate sarcomeres. However, the instructive role of titins for sarcomeric architecture outside of vertebrates is not as well understood. Here, we used a series of nanobodies, the *Drosophila* titin nanobody toolbox, recognising specific domains of the two *Drosophila* titin homologs Sallimus and Projectin to determine their precise location in intact flight muscles. By combining nanobodies with DNA-PAINT super- resolution microscopy, we found that, similar to vertebrate titin, Sallimus bridges across the flight muscle I-band, whereas Projectin is located at the beginning of the A- band. Interestingly, the ends of both proteins overlap at the I-band/A-band border, revealing a staggered organisation of the two *Drosophila* titin homologs. This architecture may help to stably anchor Sallimus at the myosin filament and hence ensure efficient force transduction during flight.

## Introduction

Skeletal and heart muscles produce forces that power body movements and fluid flow in animals. These forces are produced by conserved macromolecular machines called sarcomeres. Sarcomeres are organised into long periodic chains called myofibrils that mechanically span the entire muscle fiber length and thus sarcomere contraction results in muscle contraction (Gautel, 2011; Huxley, 1969; Lemke and Schnorrer, 2017).

The sarcomeric architecture is conserved in striated muscles across animals. Sarcomeres are bordered by two Z-discs, which anchor the plus ends of parallel actin filaments. These extend towards the centrally located bipolar myosin filaments that are cross- linked at the M-band of the sarcomere. In vertebrate sarcomeres, actin and myosin filaments are mechanically linked by the connecting filament built by the gigantic titin protein, whose N-terminus is anchored to alpha-actinin at the Z-disc while its C-terminus is embedded within the sarcomeric M-band. Thus, titin spans as linear protein across half a sarcomere in vertebrate muscle (Gautel and Djinović-Carugo, 2016; Lange et al., 2006; Linke, 2018; Squire et al., 2005). This stereotypic sarcomere architecture results in a defined sarcomere length (distance between two neighbouring Z-discs), which is about 3 µm in relaxed human skeletal muscle (Ehler and Gautel, 2008; Llewellyn et al., 2008; Regev et al., 2011), and is responsible for the typical striated appearance of skeletal muscles.

The defined sarcomeric architecture sparked the ‘titin ruler hypothesis’, proposing that the long titin protein rules sarcomere length in vertebrate muscles (Tskhovrebova and Trinick, 2012; 2017). Recently, this hypothesis has been strongly supported by *in vivo* genetic evidence. Deletion of parts of titin’s flexible I-band or its stiff A-band regions in mouse skeletal muscle resulted in a shortening of the sarcomeric I-band or A-band, respectively (Brynnel et al., 2018; Tonino et al., 2017). Furthermore, recent evidence substantiated that titin is the main sarcomeric component responsible for the passive tension of the muscle, suggesting that mechanical tension present in relaxed muscle is stretching titin into its extended conformation (Li et al., 2020; Linke, 2018; Rivas-Pardo et al., 2020; Swist et al., 2020). Thus, titin mechanically links actin and myosin filaments together and is responsible for establishing and maintaining sarcomeric architecture in vertebrate striated muscle.

Striated muscle architecture is not restricted to vertebrates but conserved in insects and worms. However, in contrast to vertebrates, titin’s role in *Drosophila* and *C. elegans* appears to be split into two proteins, one containing the flexible I-band features and the other the stiff A-band features of titin (Loreau et al., 2022; Tskhovrebova and Trinick, 2003). Surprisingly, the sarcomere length in flies and worms is still stereotypic for the respective muscle fiber type. In *Drosophila* the sarcomere length is about 3.5 µm for indirect flight muscles and about 8 µm for larval body wall muscles. To date, it is unclear how sarcomere length in these muscles is determined. Furthermore, it is unknown how the titin homologs are precisely organised within the sarcomere and if they contribute to sarcomere length regulation in insect muscle.

We aim to address the questions how invertebrate titin homologs instruct sarcomere architecture and if the titin nano-architecture would be consistent with a ruler function mechanically linking actin to myosin at a defined distance, as proposed for vertebrates. A first step to answer these important questions is to determine the exact positions of the titin homologs within the sarcomere.

Here, we chose the *Drosophila* indirect flight muscles to determine the precise location of the two *Drosophila* homologs Sallimus (Sls) and Projectin (Proj). We selected key domains at different locations within Sls and Proj, against which we raised specific nanobodies (Loreau et al., 2022). We applied single and dual-colour DNA-PAINT super- resolution microscopy to intact flight muscle specimens, which determined the precise architecture of Sls and Proj in the flight muscle sarcomere. Interestingly, we found that Sls but not Proj extends from the Z-disc to the myosin filament. The end of Sls overlaps with the beginning of Proj, which further projects along the myosin filament. This staggered organisation of the two *Drosophila* titin homologs may explain how high mechanical tension can be stably transmitted across the sarcomere and how sarcomere length can be ruled without the presence of a single protein linking the Z-disc to the M-band as observed in vertebrates.

## Results

### *Drosophila* titin domain organisation and flight muscle isoforms

*Drosophila* indirect flight muscles (called flight muscles in the remainder of the manuscript) are stiff muscles that oscillate at high frequency to power flight (Dickinson, 2006; Pringle, 1981; Schönbauer et al., 2011). The majority of this stiffness is due to Sls in flight muscles (Kulke et al., 2001). To achieve this high stiffness, a large part of the flexible spring domains encoded in both titin gene homologs *sls* and *bent* (*bt*; protein named Projectin) are spliced out by alternative splicing (Ayme-Southgate et al., 2005; Bullard et al., 2005; Burkart et al., 2007; Spletter et al., 2015). Older work had suggested that the most prominent Sls flight muscle isoform (also called Kettin) uses an alternative poly-A site terminating the protein after Sls- immunoglobulin (Ig) domain 35 (Bullard et al., 2005; Burkart et al., 2007). However, more recent systematic transcriptomics and splice-site annotation data from dissected flight muscles, as well as expression of large genomic Sls-GFP tagged transgenes, showed that usage of this early poly-A site is largely restricted to leg muscles and hardly present in flight muscles (Spletter et al., 2015; 2018). To identify the most prominent Sls and Proj protein isoforms in mature flight muscles, we carefully reanalysed the published transcriptomics and splice data (Spletter et al., 2015; 2018). We verified that in both genes, the flexible PEVK spring domains are largely spliced out in adult flight muscles, but their more C-terminally located exons are present at least in some longer isoforms (Figure 1 – figure supplement 1A, B). This predicts a Sls isoform containing the C-terminal five fibronectin (Fn) domains and a Proj isoform containing a long stretch of Ig-Fn super-repeats and a kinase domain close to its C-terminus (Figure 1 – figure supplement 1A, B).

**Figure 1.**
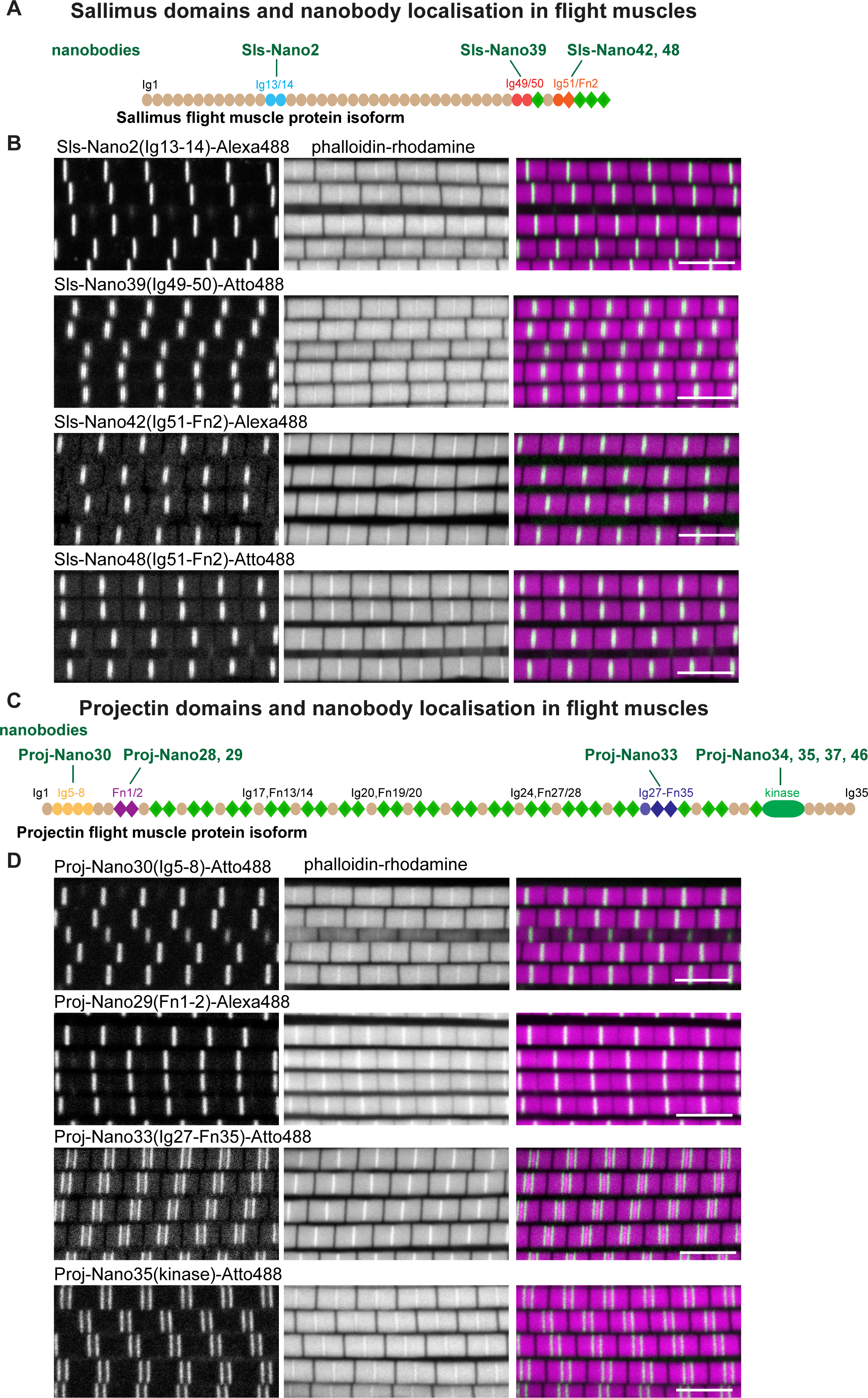
*Drosophila* titin domain organisation and nanobodies. (**A, C**) Sallimus (A) and Projectin (C) flight muscle protein isoforms with the domains recognised by the used nanobodies highlighted in different colours. (**B, D**) Single confocal sections of flight muscle sarcomeres from adult hemi-thoraces stained for actin with phalloidin (magenta) and the indicated Sls or Proj nanobodies directly coupled to Alexa488 or Atto488 (green). The Z-disc is revealed by the prominent actin signal. Scale bars 5 µm.

### Sallimus and Projectin nanobodies in flight muscles

In order to verify the expression and to determine the precise location of the different Sls domains in adult flight muscle sarcomeres, we selected three different regions in Sls, against which we recently generated nanobodies (Loreau et al., 2022): Sls-Ig13/14, Sls-Ig49/50 and Sls-Ig51/Fn2, the first being relatively close to the N-terminus, the other two being close to the C-terminus of the Sls flight muscle isoform (Figure 1 – figure supplement 1A). Similarly, we selected two regions in Proj close to its N-terminus (Proj-Ig5-8 and Proj-Fn1/2) and two regions close to its C-terminus (Proj-Ig27-Fn35 and Proj-kinase domain). The generation of these nanobodies against Sls and Proj domains as well as their specificity, tested in *Drosophila* embryonic and larval muscles, were documented in an accompanying manuscript (Loreau et al., 2022).

To verify nanobody specificity in flight muscles, we stained *Drosophila* dorso-longitudinal flight muscles with one nanobody together with phalloidin revealing the sarcomere architecture. As expected, using scanning confocal microscopy, we found Sls- Nano2 (binding Sls-Ig13/14) in a single sharp band overlapping with the sarcomeric Z-disc (Figure 1A, B). Interestingly, Sls-Nano39 (recognising Sls-Ig49/50), Sls-Nano42 and Sls- Nano48 (the latter two binding to SlsIg51/52) result in slightly broader bands at the Z-discs (Figure 1B). These data further support that the Sls nanobodies are specific and they demonstrate that the Sls C-terminal epitopes are indeed expressed in flight muscles, as predicted from the splicing data.

Similarly, we investigated the nanobodies raised against the four different Projectin domains. We found that Proj-Nano30 (recognising Proj-Ig5-8 very close to the N-term of Proj), Proj-Nano28 and Proj-Nano29 (both binding Proj-Fn1-2) each result in a broad band at the Z-disc (Figure 1C, D, Figure 1 – figure supplement 1C). Furthermore, we found that the C-terminally located Proj-Nano33 (binding Proj-Ig27-Fn35) and Proj-Nano34, Proj-Nano35, Proj-Nano37 and Proj-Nano46 (all binding the Projectin kinase domain) result in two bands at large distances from the Z-disc (Figure 1C, D, Figure 1 – figure supplement 1C). These data demonstrate that Projectin is present in an extended conformation and since the flight muscle I-band extends less than 100 nm from the Z-disc (Burkart et al., 2007; Kronert et al., 2018; Loison et al., 2018; Reedy and Beall, 1993; Szikora et al., 2020), a large part of Projectin is present along the myosin filament. However, the diffraction-limited spatial resolution of a confocal microscope is not sufficient to precisely localise Sls and Proj domains close to the Z-disc. Hence, higher spatial resolution is necessary to determine the precise architecture of Sls and Proj within the flight muscle sarcomere.

### Nanobodies have superior penetration compared to antibodies in flight muscles

Nanobodies are only 13 kDa or less than 4 nm in size (Helma et al., 2015; Pleiner et al., 2015) and hence are ideal labels for two main reasons: their small size is first placing the label very close to the domain of interest and second allows efficient penetration into dense structures present in cells or protein complexes which enables high labelling density. This was demonstrated by the superior labelling abilities of our nanobodies compared to antibodies in stage 17 embryos in an accompanying manuscript (Loreau et al., 2022). As an attempt to verify if this is also the case in the crowded environment of mature flight muscles, we stained flight muscles with Sls-Nano2 (binding Sls-Ig13/14) and compared them to the endogenously expressed M-band protein Obscurin-GFP, or to a staining with an anti-Sls antibody (anti-Kettin, binding Sls-Ig16). We imaged 10 µm thick z-stacks to quantify label diffusion into the thick and crowded flight muscle fibers. Because of light scattering and fundamental limits of confocal imaging, intensity of endogenously expressed labels also decays with imaging depth (Sarov et al., 2016). Using the same imaging conditions and same fluorophore for Sls-Nano2 and the combination of anti-Sls primary and secondary antibodies, we found that the Sls- Nano2 intensity decay over z-depth is about 2.5-fold lower than the one of the anti-Sls antibody label (Figure 2A-C, Figure 2 – figure supplement 1). This strongly suggests better penetration of the nanobody into the muscle samples compared to the larger primary and secondary antibodies.

**Figure 2.**
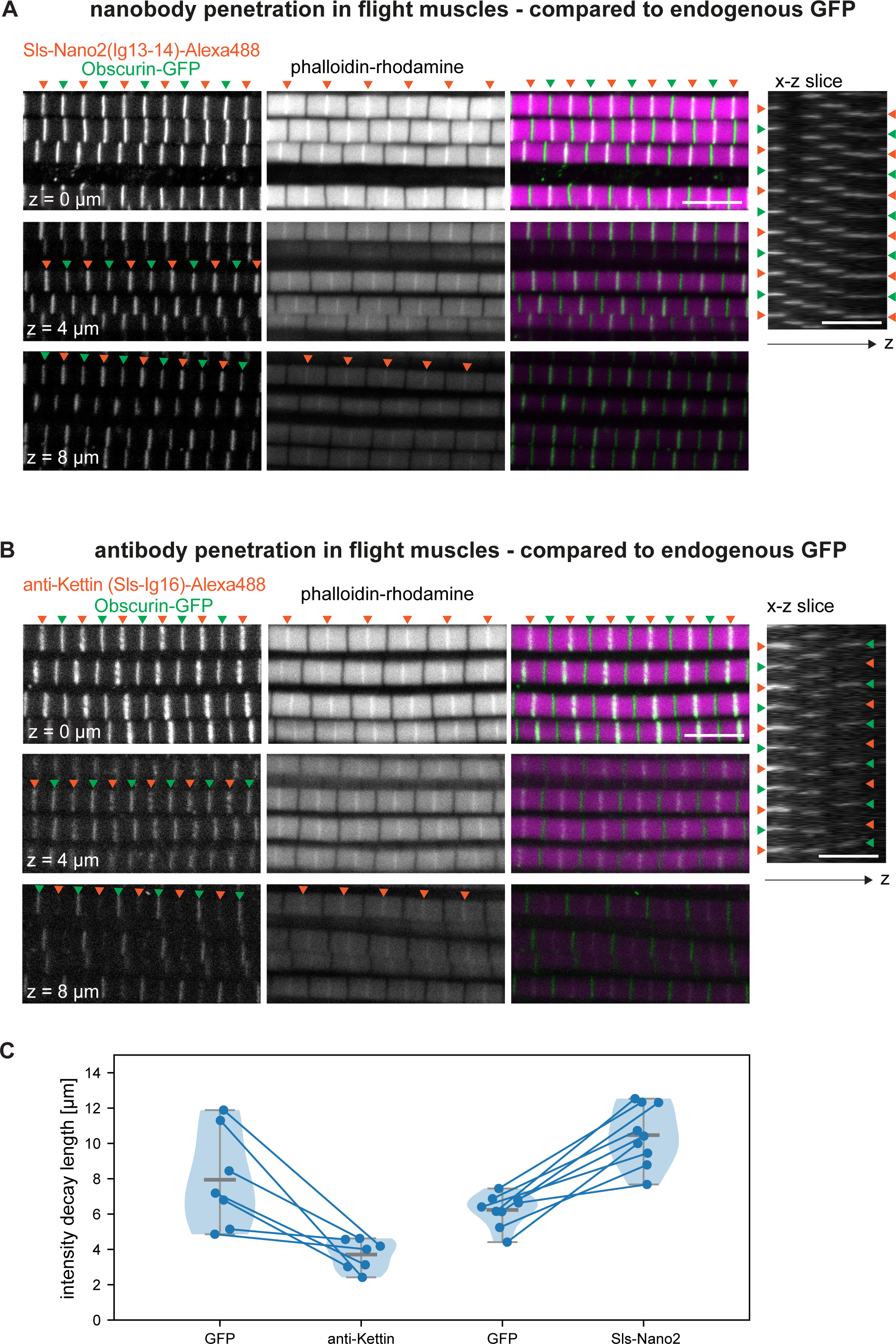
Nanobody penetration. (**A, B**) Adult hemi-thorax expressing Obscurin-GFP (green) in flight muscles stained with phalloidin to label actin (magenta) and either Sls-Nano2-Alexa488 (A) or anti-Kettin antibody (binding Sls-Ig16) (red), followed by secondary antibody coupled with Alexa488 (B). Three different z-planes and x-z slice are shown. Note that nanobody (red arrowheads in A) and GFP signals (green arrowheads) are visible in the entire z-stack, whereas the antibody signal decays quickly in z-direction (red arrowheads in B). Scale bars 5 µm. (**C**) Fluorescence detection decay length versus imaging depth for GFP, anti-Kettin and Sls-Nano2 (anti-Kettin vs Sls-Nano2 comparison: p-value = 0.0001748, Mann-Whitney test).

To directly compare the diffusion of the differently sized labels in the same samples, we double-stained flight muscles with Sls-Nano2 and the Sls antibody. We swapped the dye colours to rule out any bias of the excitation wavelength on penetration depth. We found that Sls-Nano2 readily diffuses into the thick flight muscle samples, whereas the Sls antibody is limited to the top layer of myofibrils (Figure 2 – figure supplement 2A, B). This demonstrates the favourable diffusion properties of the small nanobodies in the crowded environment of adult flight muscles. Labelling of myofibrils in the past was often achieved on isolated myofibrils to improve antibody accessibility (Burkart et al., 2007; Szikora et al., 2020), but myofibril isolation may change sarcomere mechanics and thus lead to unwanted mechanical or structural artefacts (Ayme-Southgate et al., 2004; Kulke et al., 2001). The properties of nanobodies make us confident that direct labelling of intact flight muscle fibers will lead to dense labelling with limited structural artefacts.

### DNA PAINT super-resolution imaging of entire flight muscles

In order to make use of the small size of the nanobodies, we turned our attention to super-resolution imaging with DNA-PAINT (Jungmann et al., 2014; Lelek et al.; Schnitzbauer et al., 2017). For DNA-PAINT, nanobodies binding the protein epitope of interest are site-specifically conjugated to either one or two single-stranded DNA molecules (see Methods). In DNA-PAINT, the necessary target blinking for localisation-based super-resolution reconstruction is achieved by the transient binding of dye-labelled DNA ‘imager’ strands to their target-bound complements (‘docking’ strands, Figure 3A). As imager strands are continuously replenished from solution and binding times are controllable over a wide range, a large number of photons can be detected from a single binding event, thus enabling unprecedented sub-5 nm spatial resolutions (Dai et al., 2016; Schnitzbauer et al., 2017).

**Figure 3.**
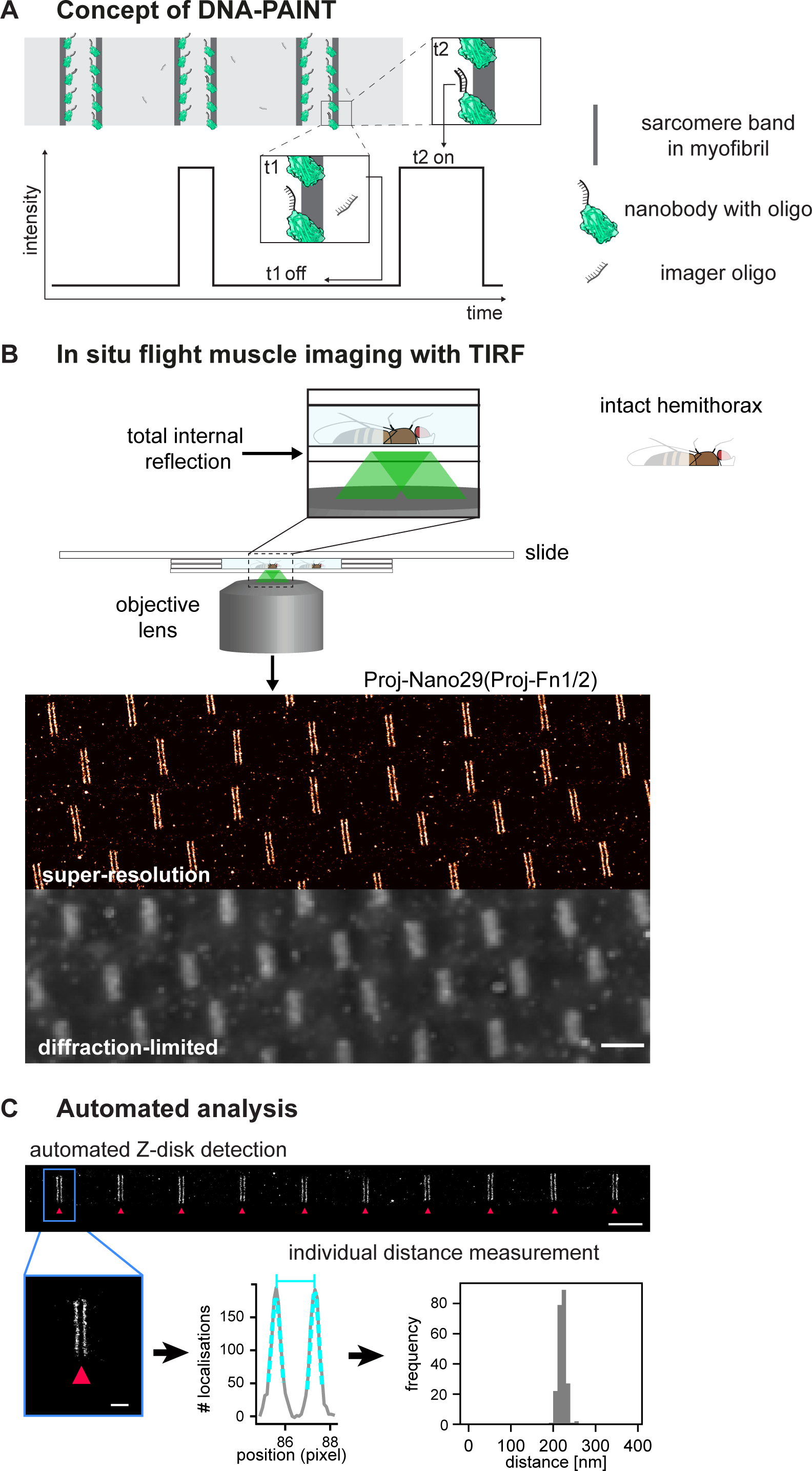
*Drosophila* flight muscle DNA-PAINT imaging and automated extraction of sarcomeric protein domain positions. (**A**) Concept of DNA-PAINT imaging of sarcomeres labelled with an oligo-conjugated nanobody. Binding of the imager oligo to one nanobody results in a strong, detectable intensity burst (t2, blink). (**B**) Schematic of a mounted intact *Drosophila* hemi-thorax in a DNA-PAINT imaging chamber enabling TIRF illumination. Comparison of the diffraction-limited and the super-resolved result illustrated in one hemi-thorax labelled with Proj-Nano29. Note that the super-resolved image can readily resolve the two-bands flanking each Z-disc. Scale bar 2 µm. (**C**) Automated image analysis for individual Z-discs detection (see Figure 3 Supplement 3 and Methods for details). Individual bands are detected automatically and their center position is obtained using a Gaussian fit (bottom center). The distance between the center of bands for tens of sarcomeres from a single hemi-thorax is then reported in a histogram (bottom right). Scale bar 2 µm (top) and 0.5 µm (bottom).

Previously, DNA-oligos for PAINT were either coupled to antibodies via biotin-streptavidin (Jungmann et al., 2014), which is a 66 kDa tetramer and thus relatively large or more frequently by click chemistry (Fabricius et al., 2018; Schnitzbauer et al., 2017), which comes with a number of potential disadvantages, such as a bulky hydrophobic coupling group and an initial lysine modification that might destroy the paratope. Instead, we used maleimide-coupling through cysteines at the N- and C-terminus of the nanobody, which allows a simpler workflow, analogous to direct fluorophore coupling, and protects the antigen-binding site from undesired modifications.

In contrast to fluorophore-maleimides, maleimide-activated oligonucleotides are not commercially available. However, as described in the Methods, they are straightforward to synthesise from a 5’ amino-modified oligo and a bifunctional maleimide-NHS (N-hydroxysuccinimide) crosslinker. The NHS group forms an amide bond with the 5’ amino group of the oligo under reaction conditions that leave the amino groups of the DNA bases non-reactive. The maleimide-activated oligo is then reacted with the nanobody that still contains its His14-SUMO or His14-NEDD8 tag. The resulting conjugate is purified by binding to a Ni(II) chelate matrix (whereby the non-conjugated excess of oligo remains in the non-bound fraction) and followed by elution of nanobody-oligo conjugate with a tag-cleaving protease. Hence, these oligo-coupled nanobodies remain similarly small as the fluorescently coupled nanobodies and are thus ideal for effective super-resolution imaging using DNA-PAINT.

We aimed to apply DNA-PAINT to flight muscle tissue, using hemi-thoraces of adult flies, to minimise artefacts that might be introduced by cutting out individual myofibrils. To prepare hemi-thoraces, we fixed thoraces in paraformaldehyde and then bisected them with a sharp microtome knife (Figure 3 – figure supplement 1A, see Methods for details). Then, we incubated the hemi-thoraces with oligo-coupled nanobodies and mounted them for imaging. Hemi-thoraces are very large, with a length of about 1 mm and a thickness of about 300 µm. To mount them as close as possible to the coverslip, we developed an imaging chamber that contains the imaging buffer surrounded by spacers thick enough to slightly press the flight muscles against the coverslip (Figure 3 – figure supplement 1A, see Methods for details). This enabled DNA-PAINT imaging with total internal reflection (TIRF). We imaged for 30 min per sample and obtained about 15000 frames at an imaging rate of 10 Hz. For image reconstruction and post-processing, we used the established Picasso software (Schnitzbauer et al., 2017) (Figure 3B, Figure 3 – figure supplement 1B, see Methods for details). This enabled us to resolve the two bands flanking a Z-disc with ease, which cannot be resolved in the diffraction-limited image (Figure 3B).

To further refine the precision of determining the epitope positions, we have developed an image processing pipeline that relies on an interactive selection of well-stained myofibrils in the volume of TIRF excitation (Figure 3 – figure supplement 2). Next, we removed localisations arising from multiple binding events by filtering based on specific localisation parameters (see Methods for details). Further, we automatically detected the individual sarcomeric Z-discs and the respective flanking bands of the stained Sls or Proj epitopes for all selected myofibrils. We applied a Gaussian fit to each band and determined their center positions within the sarcomere with nanometric accuracy (Figure 3C, Figure 3 – figure supplement 2). This results in an accurate location of the measured bands for each of the epitopes in every analysed sarcomere. Hence, we do not need to average across many sarcomeres to precisely localise the Sls or Proj epitopes (Figure 3C). In conclusion, our method allows detecting individual differences in sarcomeric band positions in each sarcomere investigated down to the nanometer-scale.

### Positions of Sallimus and Projectin domains within intact flight muscle at the nanometric scale

We applied our DNA-PAINT imaging pipeline of flight muscles to the Sls and Proj nanobody toolbox. In most cases, we co-stained with two nanobodies that are spaced sufficiently apart to detect the expected four bands centred around the Z-disc, even when using only a single imaging colour (Figure 4). This allowed us to resolve the positions of Sls-Nano2 (Sls-Ig13/14) located close to the N-terminus of Sallimus or Sls-Nano39 (Sls-Ig49/50) close to its C-terminus, which we could combine with distantly located Proj nanobodies (Figure 4A). Similarly, we imaged the N-terminally located Proj-Nano29 (Proj-Fn1/2) and Proj-Nano30 (Proj-Ig5-8), which we could combine with one of the C-terminally located Proj-Nano33 (Ig27-Fn35), Proj-Nano35 or Proj-Nano37 (both Proj kinase domain). This enabled us to locate the exact position of different Projectin domains in sarcomeres (Figure 4A). Interestingly, all the analysed epitopes result in similarly sharp bands in each of the sarcomeres, suggesting a very precise linear architecture of Sls and Proj. The N-terminus of Sls is located close to the Z-disc, with Ig13/14 only about 50 nm away from the center of the Z-disc, whereas the N-terminus of Proj is located around 100 nm away from the Z-disc (Proj- Ig5-8 and Proj-Fn1/2) and hence cannot be anchored directly at the Z-disc.

**Figure 4.**
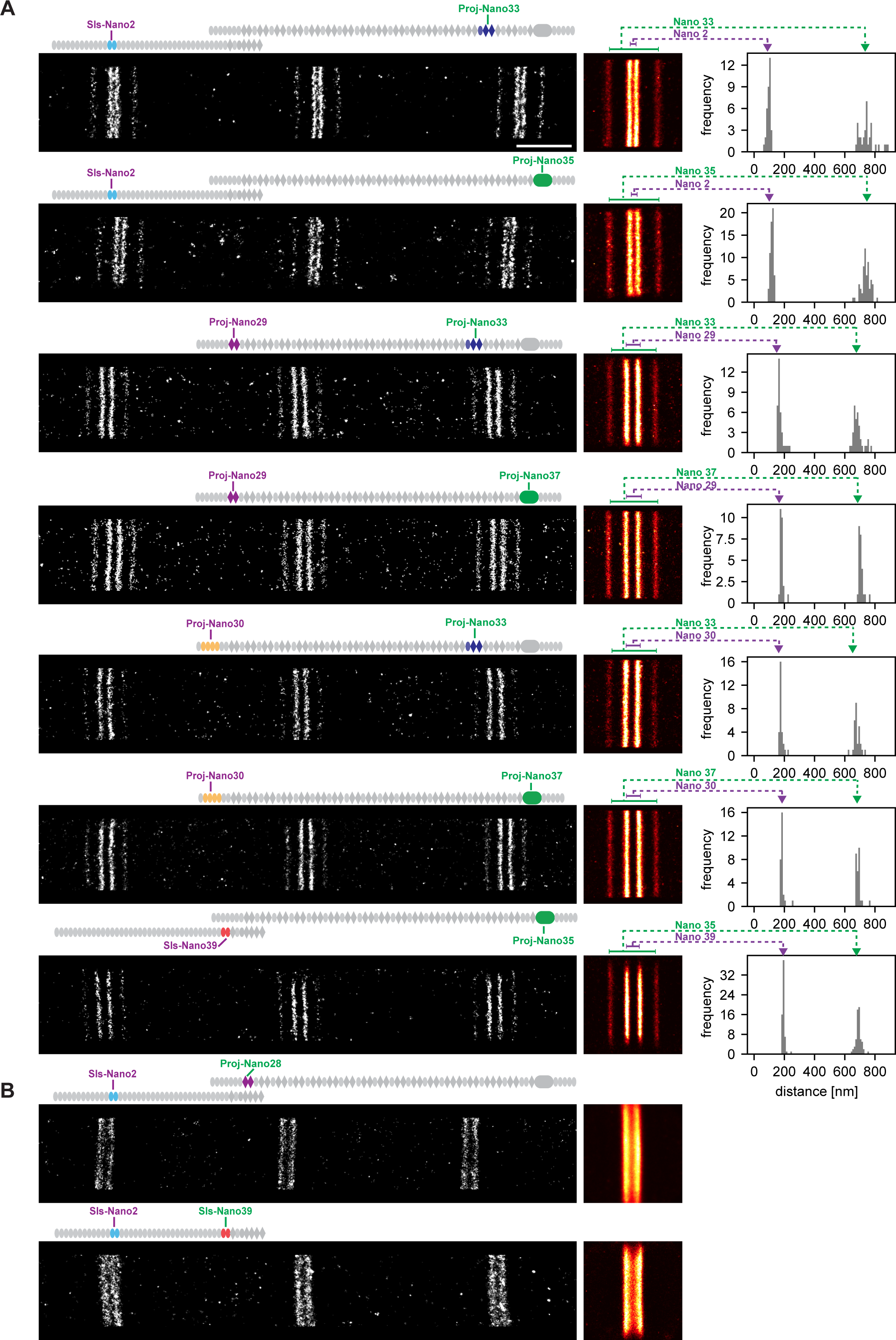
Single-colour DNA-PAINT imaging of Sls and Proj domains. (**A**) Left: representative DNA-PAINT images of myofibrils stained with two different Sls or Proj nanobodies labelling two epitopes and imaged with the same fluorescent imager oligo. The different Sls or Proj nanobody combinations are indicated above each image. Middle: pseudo-coloured sum image centered around Z-discs resulting from one hemi-thorax. Right: histogram of distances between bands centered around Z-discs with the respective nanobody combinations indicated in green or magenta. Note that four bands can be readily distinguished for all shown nanobody combinations. (**B**) Similar representations as in (A). However, the positions of neighbouring Sls or Pro epitopes cannot be resolved in a single colour. Scale bar 1 µm.

This single-colour imaging method is powerful, but it fails to resolve two epitopes into distinct bands if the epitopes are located too close together to unambiguously assign each blinking event to one particular nanobody. Thus, Sls-Nano2 (Sls-Ig13/14) and Sls-Nano39 (Sls-Ig49/50) or Sls-Nano2 (Sls-Ig13/14) and Proj-Nano28 (Proj-Fn1/2) cannot be imaged together in the same sarcomere with a single colour (Figure 4B). However, quantifying the exact positions of two closely located titin domains in the same sarcomere is critical as the relative length of the flexible titin molecules may vary in individual sarcomeres. Hence, it would be important to determine the positions of two different Sls domains in the same sarcomere to unambiguously conclude about Sls length or the relative arrangement of its protein domains.

### Two-colour DNA-PAINT reveals a staggered organisation of Sls and Proj

To simultaneously determine the exact positions of two epitopes, we have labelled our nanobodies with two different oligonucleotides and imaged them with two different imager oligos in parallel to perform two-colour DNA-PAINT (see Methods). Multiplexed imaging enabled us to determine the positions of Sls-Ig13/14 (using Sls-Nano2) and Sls-Ig51/Fn2 (using Sls-Nano42) in the same sarcomere (Figure 5A). Our results verified that Sls-Ig13/14 is localised about 50 nm away from the center of the Z-disc and that Sls-Ig51/Fn2 is about 50 nm further towards the middle of the sarcomere (Figure 5 – figure supplement 1). Since the I-band of flight muscles is less than 100 nm from the Z-disc (Burkart et al., 2007; Kronert et al., 2018; Loison et al., 2018; Reedy and Beall, 1993; Szikora et al., 2020) this strongly suggests that Sls is bridging across the entire sarcomeric I-band with its N-terminus anchored within the Z-disc and its C-terminal end reaching the myosin filament. Thus, Sls could mechanically link the Z-disc to the myosin filament in the flight muscles, similar to the long vertebrate titin.

**Figure 5.**
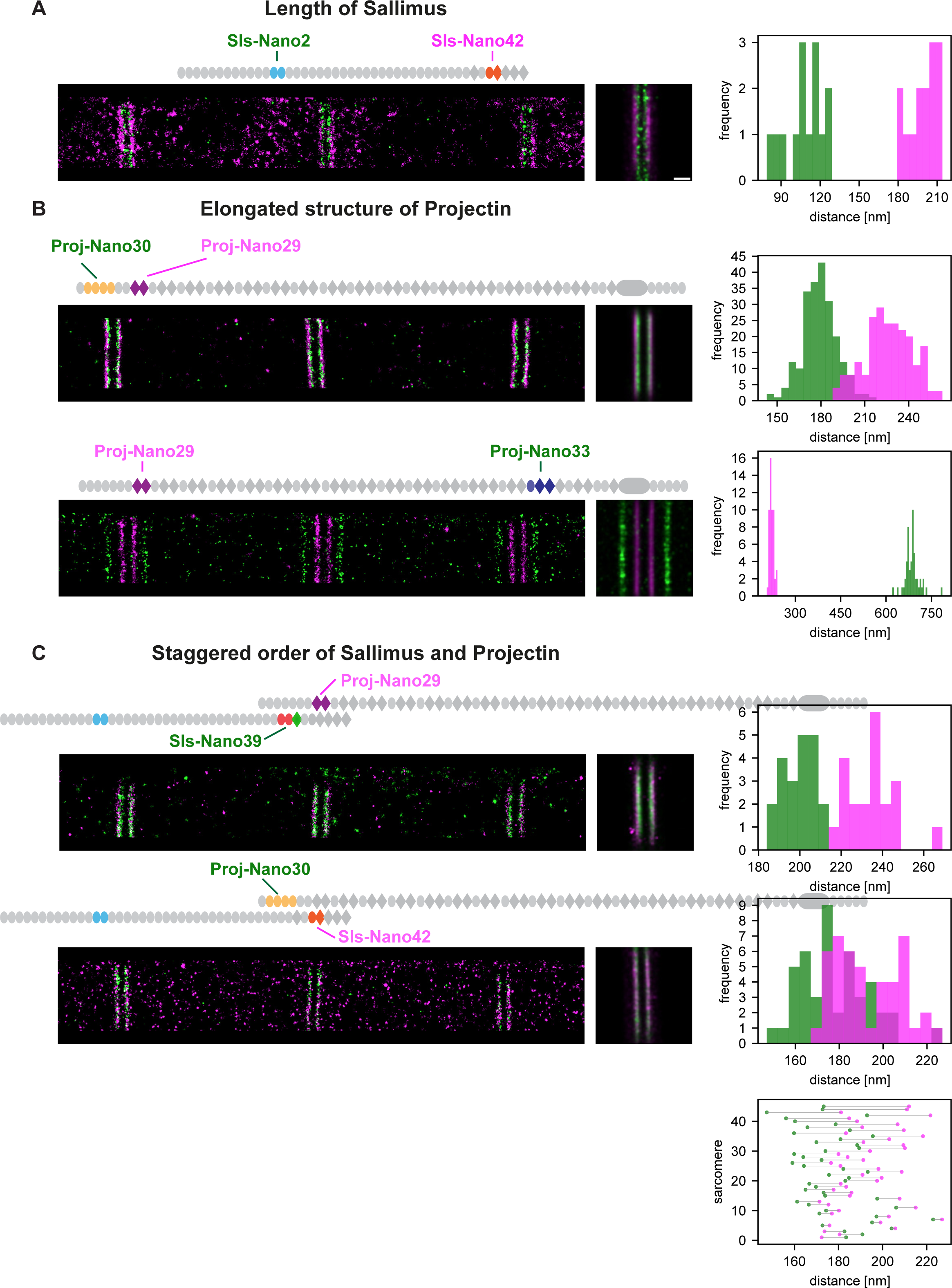
Dual-colour DNA-PAINT imaging reveals staggered order of Sls and Proj. (**A**) Left: representative DNA-PAINT image of a myofibril stained with two nanobodies labelling Sls-Ig13/14 (Sls-Nano2) and Sls-Ig51/Fn2 epitopes (Sls-Nano42). Middle: sum image centered around Z-discs resulting from one hemi-thorax. Right: histogram of distances between bands centered around Z-discs (Sls-Ig13/14 in green, Sls-Ig51/Fn2 in magenta). (**B**). Top: representative DNA-PAINT image of a myofibril stained with two nanobodies labelling Proj-Ig5-8 (Proj-Nano30) and Proj-Fn1/2 (Proj-Nano29) epitopes, sum image and histograms of distances between bands (Proj-Ig5-8 in green, Proj-Fn1-2 in magenta). Bottom: representative myofibril stained for Proj-Fn1/2 (Proj-Nano29) and Proj-Ig27-Fn35 (Proj-Nano33) epitopes, sum image and histogram of distances between bands centered around Z-discs (Proj-Fn1/2 in magenta, Proj-Ig27-Fn35 in green) (**C**). Top: representative DNA-PAINT image of a myofibril stained with two nanobodies labelling SlsIg49/50 (Sls-Nano39) and Proj-Fn1/2 (Proj-Nano29) epitopes, sum image and histogram of distances between bands centered around Z-discs (Sls-Ig49/50 in green, Proj-Fn1/2 in magenta). Bottom: same as top for Sls-Ig51/Fn2 (Sls-Nano42) and Proj-Ig5-8 (Proj-Nano30) epitopes, sum image, histogram of distances and plot showing the epitope positions from the Z-discs in the individual sarcomeres analysed (bottom right, Sls-Ig51/Fn2 in magenta, Proj-Ig5-8 in green). Note that in 42 of 45 cases the Proj-Ig5-8 (green) is closer to the Z-disc than Sls-Ig51/Fn2 (magenta). Scale bar 250 nm.

Since we found that the N-terminus of Proj is also about 100 nm away from the Z-disc and thus located at the beginning of the thick filament (Figure 4A) we wanted to further investigate the precise orientation of the Proj N-terminal domains. We performed two-colour DNA-PAINT to localise Proj-Ig5/8 (with Proj-Nano30) and Pro-Fn1/2 (with Proj-Nano29) in the same sarcomere and found an average distance between the two epitopes of about 25 nm, with Proj-Ig5-8 being always closer to the Z-disc relative to Proj-Fn1/2 (Figure 5B, Figure 5 – figure supplement 1). Consistently, the more C-terminally located Proj-Ig27-Fn35 epitope is located far into the myosin filament beginning at 100 nm (Szikora et al., 2020), being 350 nm away from the Z-disc (Figure 5B, Figure 5 – figure supplement 1). This strongly suggests that the N-terminal part of Proj is arranged in an extended, likely linear confirmation reaching from the myosin filament into the I-band and thus running in parallel to the C-terminal domains of Sls.

These findings raised an enticing hypothesis: do the extended Sls and Proj proteins overlap at the I-band/A-band interface? To investigate this hypothesis, we performed two-colour DNA-PAINT using two pairs of nanobodies: Sls-Nano39, recognising Sls-49/50, combined with Proj-Nano29, binding Proj-Fn1/2 and Sls-Nano42, recognising Sls-Ig51/Fn2, combined with Proj-Nano30, binding Proj-Ig5/8. Interestingly, we found that in all sarcomeres measured, the Proj-Nano29 is about 15 nm further from the Z-disc than Sls- Nano39, whereas in 42 out of 45 sarcomeres investigated, Proj-Nano30 is on average 7-8 nm closer to the Z-disc than Sls-Nano42 (Figure 5C, Figure 5 – figure supplement 1). Hence, these data revealed an interesting staggered organisation of the two overlapping ends of the linearly extended Sallimus and Projectin proteins in flight muscles.

### A molecular map of the *Drosophila* titin homologs in flight muscle sarcomeres

Our data enabled us to build a molecular map of the *Drosophila* titin homologs in flight muscle sarcomeres, which revealed a significant overlap of the linear Sls and Proj proteins at the I-band/A-band interface as visualised in a ‘composite sarcomere’ reconstructed by imaging flight muscles from six different hemi-thoraces (Figure 6A).

**Figure 6.**
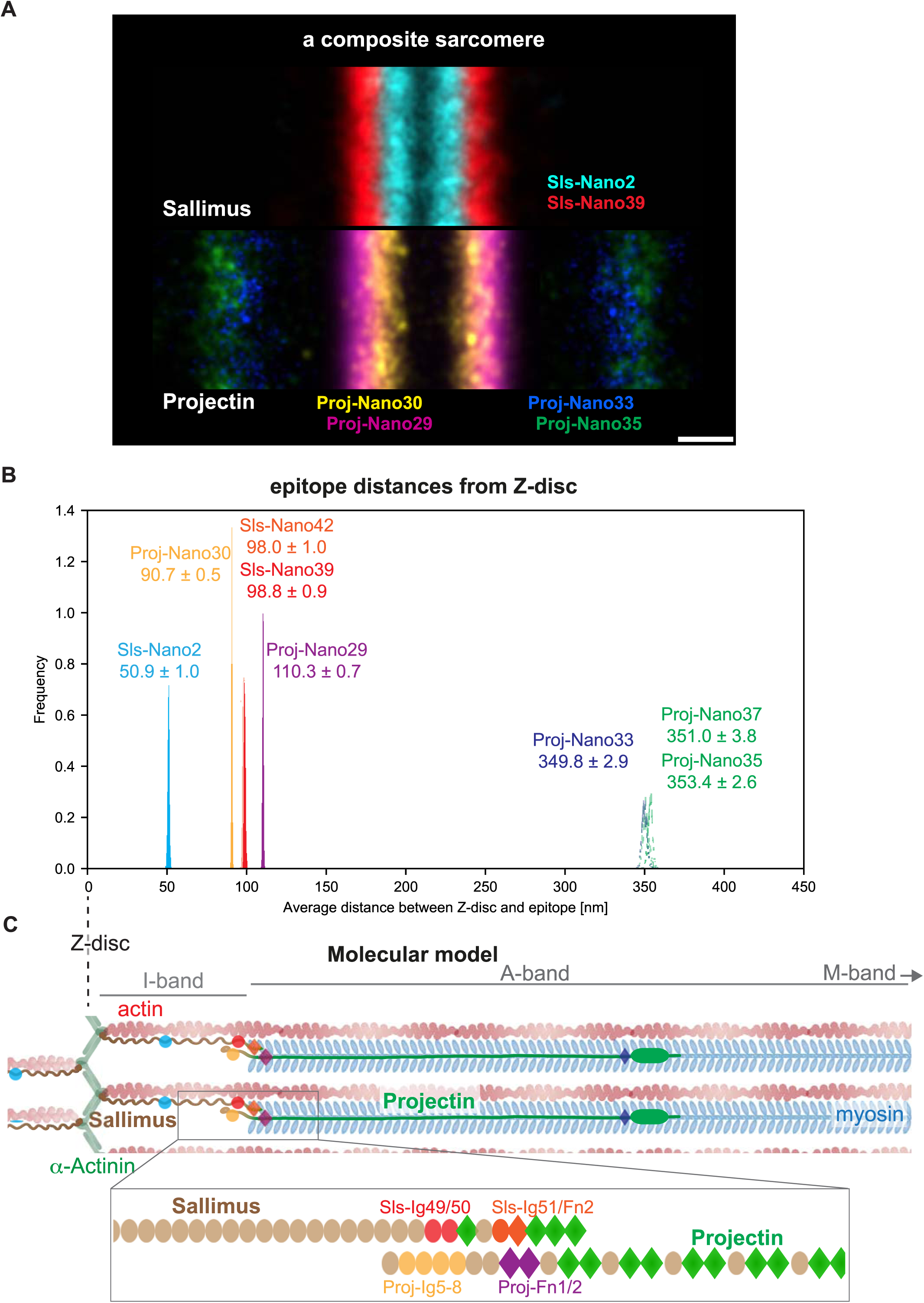
Summary and model. (**A**) A sarcomere displayed as a composite of two summed Sls nanobody bands (top) and four summed Proj nanobody bands (bottom), each originating from one individual hemi-thorax imaged. Note the overlay of the positions of both proteins. Scale bar is 200 nm. (**B**) Distribution of the averaged distances from the Z-disc for all Sls and Proj epitopes measured using bootstrapping (see Methods). (C) Cartoon model of the relative arrangement of Sls and Proj within the flight muscle sarcomere. The positions of the measured Sls and Proj domains are highlighted in colours. The zoomed regions illustrate the suggested staggered architecture of the C-terminal Sls and the N-terminal Proj protein parts.

To precisely determine the position of all the epitopes investigated in our study, we calculated the averaged position using all the sarcomeres we imaged in the single- and dual-colour DNA-PAINT experiments. This strategy is valid as we found that although our mounting protocol for TIRF imaging results in a slightly variable sarcomere length around 3.5 µm (Spletter et al., 2015), the distance between the measured epitopes is constant (Figure 6 – figure supplement 1). Hence, the localisation of the Sls and Proj domains investigated using all sarcomeres measured resulted in a very high localisation precision with 95% confidence intervals of only 1 to 8 nm (Figure 6B). Pooling all data verified that the N-terminal Proj-Ig5-8 epitope is located 90 nm from the Z-disc, whereas the C-terminal Sls epitopes Sls-Ig49/50 and Sls-Ig51/Fn2 are located about 98 nm from the Z-disc. This is consistent with a staggered linear organisation of Sallimus and Projectin, which suggests an attractive mechanism how to mechanically link the sarcomeric Z-disc in insect flight muscle with the myosin filament using both titin homologs (Figure 6C).

## Discussion

### Super-resolution of flight muscles with nanobodies

The value of nanobodies and other small binders is well appreciated (Harmansa and Affolter, 2018). However, most *Drosophila in vivo* studies have thus far heavily relied on commercially available anti-GFP nanobodies to enhance GFP fluorescence signal in various tissues, including *Drosophila* flight muscles (Kaya-Copur et al., 2021) or to either trap GFP-fusion proteins ectopically or to degrade them when expressed in various modified forms *in vivo* (Caussinus et al., 2011; Harmansa et al., 2015; Nagarkar-Jaiswal et al., 2015). Our titin nanobody toolbox (Loreau et al., 2022) enabled us now to apply DNA-PAINT super-resolution technology to image the titin nanostructure in large intact flight muscle tissue at nanometer scale resolution.

It had been shown that dye- or DNA-labelled nanobodies work well to achieve high labelling densities in cell culture (Agasti et al., 2017; Fabricius et al., 2018; Mikhaylova et al., 2015; Pleiner et al., 2015; Schlichthaerle et al., 2019). We show here that our nanobodies are also very efficient in penetrating into the large flight muscle fibers containing highly packed sarcomeres, which are amongst the most protein dense macromolecular structures in biology (Daneshparvar et al., 2020; Taylor et al., 2019). This enabled us to perform DNA-PAINT super-resolution microscopy of the large flight muscles without dissecting individual myofibrils. Such large specimen have rarely been investigated with DNA-PAINT (Cheng et al., 2021; Lelek et al. 2021). This shows that DNA-PAINT can be readily applied to super-resolve structures in large tissues if mounting and labelling protocols are optimised.

### Titin nanoarchitecture in flight muscles – do titins rule?

Flight muscles are an ideal tissue to perform architectural studies of their sarcomeric components at the nanoscale, because these components display an extremely high molecular order (Loison et al., 2018). This was impressively demonstrated by substructural averaging that resolved the nanostructure of myosin filaments isolated from insect flight muscles at a 7Å resolution by cryo-electron-microscopy (Daneshparvar et al., 2020; Hu et al., 2016). Another recent study took advantage of this stereotypic order and used a series of existing antibodies against sarcomeric protein components to probe isolated myofibrils from *Drosophila* flight muscles using STORM (Rust et al., 2006). The high order enabled averaging of several hundred sarcomeres to reconstruct distances of the epitopes from the Z-disc with 5-10 nm precision (Szikora et al., 2020). Although done on isolated myofibrils, the large diversity of antibodies studied gave a comprehensive understanding of domain positions for a variety of important sarcomeric components. This included the Sls-Ig16 antibody used here, locating Sls-Ig16 about 50 nm from the center of the Z-disc (Szikora et al., 2020), which is in good agreement with the location of Sls-Ig13/14 we found here. This study further showed that the important Z-disc components alpha-Actinin, Zasp52 and Filamin extend only about 35 nm from the center of the Z-disc (Szikora et al., 2020). This strongly suggests that the N-terminus of Sls, with its remaining 12 Ig domains can reach and interact with these Z-disc components, as has been reported biochemically (González-Morales et al., 2017; Liao et al., 2016). Hence the N-terminal part of the fly titin homolog Sls is arranged similarly to the N-terminus of vertebrate titin that binds to alpha-Actinin, anchoring it within the Z-disc (Gautel and Djinović-Carugo, 2016; Ribeiro et al., 2014).

An important part of the titin ruler model is that the titin spring part, which relaxes and stretches during muscle contraction and relaxation, respectively, spans across the I-band and sets the I-band length of vertebrate sarcomeres (Brynnel et al., 2018; Linke, 2018; Luis and Schnorrer, 2021). Thus, it is insightful that our newly developed C-terminal Sls nanobodies show that the C-terminal end of Sls is located about 100 nm from the center of the Z-disc in flight muscles. Although we have not imaged myosin directly in our samples, both STORM and electron-microscopy studies demonstrated that the myosin filament begins about 100 nm from the center of the Z-disc, making the I-band less than 100 nm wide (Burkart et al., 2007; Kronert et al., 2018; Loison et al., 2018; Reedy and Beall, 1993; Szikora et al., 2020). This strongly suggests that, as in vertebrates, Sls is indeed spanning across the short flight muscle I-band, where it could interact with its C-terminal fibronectin domains with the myosin filament and hence could mechanically link the Z-disc with the myosin filament. This would be consistent with Sls functioning as a I-band ruler in insect muscles (see model in Figure 6C). This interpretation is also supported by the observation that non flight muscles like leg, jump and larval muscles, which contain long I-bands, do express longer versions of Sls that include the large and flexible PEVK domains (see Figure 1 Supplement 2A) (Burkart et al., 2007; Spletter et al., 2015). Indeed, in the accompanying paper using the Sls nanobodies, we showed that Sls is more than 2 µm long in larval muscles to bridge over these long I-bands (Loreau et al., 2022). This strongly suggests that Sls determines I-band length in the different muscle types, however, a direct genetic test that modifies Sls length and assays I-band length remains to be done.

The vertebrate A-band contains the Ig-Fn super-repeats of titin, which extend from the beginning myosin filament until the M-band, where titin’s C-terminal kinase is located (Granzier et al., 2014; Lange et al., 2005; Linke, 2018). Interestingly, we demonstrate that in *Drosophila* flight muscles Projectin, which is very similar to the A-band part of vertebrate titin, with long Ig-Fn super-repeats and a C-terminal kinase domain, starts about 90 nm from the Z-disc. Hence, it is very unlikely that it can interact with Z-disc components directly as these are far from the N-terminal end of Projectin (model in Figure 6C). Our precise distance measurements suggest that the N-terminus of Projectin, which does contain a series of Ig domains, typical for the I-band part of titin, is sticking into the flight muscle I-band, whereas its first Fn/Ig super-repeat is located at beginning of the A-band (110 nm from the Z-disc) and hence can interact with myosin, as can its remaining Ig-Fn super-repeats that extend over a length of about 250 nm towards the M-band. The Proj kinase localises in a sharp band, however it remains far from the M-band. Hence, it is hard to imagine that Proj alone can directly rule A-band length of flight muscle sarcomeres, as it is only present at its distal ends, spanning about 15% of the myosin filament.

### Staggering insect titins to effectively transduce forces during flight?

*Drosophila* flight muscles are very stiff to effectively power wing oscillations during flight at 200 Hz. The perpendicular arrangement of the antagonistic dorso-ventral (DVMs) versus the dorso-longitudinal flight muscles (DLMs) enables an effective stretch-activation mechanism as trigger: contraction of the DVMs moves the wings up and stretches the DLMs to induce their contraction, which will move the wings down again for the next cycle (Dickinson et al., 2005; Pringle, 1981; Syme and JOSEPHSON, 2002). The importance of strain is these muscles is highlighted by their expression of a particular troponin C isoform (TpnC4), which requires to be stretched to displace tropomyosin from myosin bindings sites on actin filaments (Agianian et al., 2004). Furthermore, myosin also experiences a stretch-induced deformation before effective actin binding and maximum force production (Iwamoto and Yagi, 2013). This strongly suggests that a very effective force transmission is needed during flight muscle oscillations.

*Drosophila* sarcomeres have a peak to peak amplitude of about 3.5 % or 60 nm per half sarcomere during flight (measured in *Drosophila virilis* (Chan and Dickinson, 1996)). This 3.5 % strain is needed to produce the up to 110 W/kg power output of insect flight muscles (Chan and Dickinson, 1996), which is consistent with the hypothesis that strain across molecules stores the elastic energy for the next contraction cycle in *Drosophila* (Dickinson et al., 2005). A perfect candidate for such a molecule is Sls as it bridges across the I-band, which likely changes length during the fast contraction cycles. Thus, Sls length would oscillate during flight, which likely results in high oscillating forces across Sls during flight.

A similar storage of elastic energy has been suggested for mammalian titin during sarcomere contraction cycles (Eckels et al., 2019; Rivas-Pardo et al., 2020).

What is the role of Projectin? The precise linear arrangement of Proj at the beginning of the myosin filament and the found overlap with Sls suggests that this staggered architecture of Sls and Proj might be required to effectively anchor Sls to the myosin filament and to prevent sarcomere rupturing during flight. Such flight induced muscle ruptures are generated when muscle attachment to tendons is weakened, underscoring the high muscle forces and high strain produced during flight (Lemke et al., 2019). Projectin may thus serve as an effective glue to stably connect Sls to the myosin filament. This is also consistent with the findings that both Sls and Proj are needed to assembly contractile sarcomeres in *Drosophila* larval muscles. Knock-down of each of the proteins results in embryonic lethality and defective sarcomerogenesis (Loreau et al., 2022; Schnorrer et al., 2010). Taken together, the staggered architecture of the two *Drosophila* titin homologs may effectively allow force transduction and ensure mechanical integrity of flight muscles sarcomeres, both very prominent functions of mammalian titin (Li et al., 2020; Rivas-Pardo et al., 2020; Swist et al., 2020).

## Methods

### Fly strains and fly culture

Fly stocks were grown and maintained under normal culture conditions in humidified incubators with 12-hour light-dark cycles on standard fly medium (Avellaneda et al., 2021). The particularly well flying ‘Luminy’ strain was used in all experiments as wild type (Leonte et al., 2021). The Obscurin-GFP strain contains a large genomic clone expressing Obscurin (unc-89) tagged with GFP at its C-terminus (Sarov et al., 2016). For all experiments young 3 to 10 day old flies were used.

### Nanobody production and labelling

Nanobody production and labelling with fluorophores by maleimide chemistry through ectopic cysteines was done as described in detail in the accompanying paper (Loreau et al., 2022). To couple nanobodies to DNA oligos, the oligos (P1, P2, PS3) were ordered with a 5’ amino group modification (e.g., Am-C6-TTT CTT CAT TAC) from IBA (Göttingen) in HPLC-purified form and lyophilized as a three ethyl ammonium (TEA) salt. Note that the absence of ammonia (NH_4_^+^) is essential for the procedure. 1 µmol of oligo was dissolved in 200 µl 30% acetonitrile (ACN), 15 mM TEA, which yielded a 5 mM stock at neutral pH (∼7). 5 µl of a 100 mM crosslinker stock in 100% ACN (maleimido β-alanine NHS ester, Iris Biotec # MAA1020 or mal-PEG4-NHS, Iris Biotec # PEG1575) were added and allowed to react for 30 minutes on ice. Then, 1.6 µl 5 M sodium acetate, and 0.1 M acetic acid (pH ∼7) were added, and the modified oligo was precipitated by adding 1 ml 100% ACN and centrifugation for 10 minutes at 0°C at 12000 rpm. This step removes any non-reacted maleimide. The pellet was then dissolved in 100µl 30% ACN, and either stored in small aliquots at -80°C or used directly to label nanobodies at ectopic, reduced cysteines, as described for fluorophores in the accompanying paper (Loreau et al., 2022). Note that free oligo cannot be removed by gelfiltration on Sephadex G25 because it appears with the conjugate in the void volume. It is best removed by modifying a still His14-SUMO, or His14- NEDD8 tagged nanobody and then using Ni(II) capture (where the free oligo remains non-bound) and proteolytic release of the then tag-free nanobody conjugate. The efficiency of conjugation can be assessed by SDS-PAGE, in which the oligo-modification results in a clear size shift. In addition, the density of modification can be calculated through OD260 and OD280 readings, using ε260 and ε280 of the initial oligo and nanobody as input variables. The oligo modification by this method is usually quantitative already with a small (≥1.1) molar excess of the maleimide oligo over modifiable cysteines. In case of incomplete modification, the conjugate can be purified on a MonoQ column, whereby the highly negative charged oligo causes stronger retention of the conjugate as compared to the non-modified nanobody.

### Flight muscle preparation, staining and mounting for imaging

Intact hemi-thoraces from adult males were prepared similar as described (Weitkunat and Schnorrer, 2014). Head, wings and abdomen were clipped with sharp forceps and the intact thoraces were fixed for 20 min at room temperature in relaxing solution (4% PFA in 100mM NaCl, 20mM NaPi pH7.2, 6mM MgCl2, 5mM ATP, 0.5% Triton X-100). After washing twice with relaxing solution, the thoraces were placed on a slide with double-sticky tape and cut sagittally with a sharp microtome blade (Pfm Medical Feather C35). The fixed hemi-thoraces were transferred to 24-well plates or Eppendorf tubes and blocked for 30 min at room temperature with 3% normal goat serum in PBS + 0.5% Tx-100 (PBS-T). Hemi-thoraces were stained overnight at 4°C with the combinations of nanobodies indicated, labelled with fluorophores or oligonucleotides (final concentration of about 50 nM). The rat anti-Sls antibody (anti-Kettin, MAC155/Klg16, Babraham Bioscience Technologies) was diluted 1:1000 and visualised with fluorescently labelled secondary antibodies (ThermoFisher, Molecular Probes). Actin was stained with phalloidin-rhodamine or phalloidin-Alexa488 (1:2000, ThermoFisher; 2 h at room temperature or overnight at 4°C). To mount the flight muscles as close as possible to the coverslip, an imaging chamber was build using a slide and #1 coverslips as spacers right and left of the samples. A layer of double sticky tape was built on the spacer and the imaging chamber was filled with either SlowFadeTM Gold Antifade (Thermofisher) for confocal imaging or Imager solution for DNA-PAINT imaging. Stained hemi-thoraces were added, oriented with the flight muscles facing up and #1.5 coverslip was added. The chamber was sealed with nail polish for confocal imaging or Picodent glue for DNA-PAINT imaging.

### Confocal imaging and processing

Stained flight muscles were imaged on a Zeiss LSM880 confocal with a 63x oil lens. Images were processed using Fiji (Schindelin et al., 2012).

### Analysing antibody versus nanobody labelling intensity decay over depth

We manually drew selections with Fiji (Schindelin et al., 2012) on stacks obtained with confocal imaging; each selection consisted of one myofibril. We used these selections to extract intensity profiles that were then analysed automatically using Python custom codes. The automated analysis to extract the intensity of each band consisted of the following: (a) locate bands in profiles using the peak finding algorithm find_peaks from the scipy library; (b) subtract background on the profile, linear fitting the 35% lowest values of the profile and subtracting this fit on the profile; (c) fit bands on the background-corrected profile with Gaussian functions; (d) estimate the area under the curve of these fits. This initial analysis allowed us to estimate the integrated intensity of bands of Obscurin-GFP and epitopes labelled with Sls-Ig16 antibody and Sls-Ig13/14 nanobody. In order to estimate how fast intensity decays with depth when imaging these bands with confocal microscopy, for each animal we fitted with an exponential decay function to the averaged band intensity over each selection (a myofibril) versus the depth where it was imaged (Figure 2 – figure supplement 1). The decay lengths obtained were then reported in Figure 2C. In our imaging conditions, the decay of intensity with depth of GFP was higher than the one of Sls-Nano2, likely caused by faster bleaching of GFP compared to the Alexa488 dye when acquiring a z-stack.

### DNA-PAINT imaging

#### Materials

Cy3B-modified and Atto643-modified DNA oligonucleotides were custom-ordered from Metabion. Sodium chloride 5 M (cat: AM9759) was obtained from Ambion. Coverslips (cat: 0107032) and glass slides (cat: 10756991) were purchased from Marienfeld and Thermo Fisher. Double-sided tape (cat: 665D) was ordered from Scotch. Two component silica twinsil speed 22 (cat. 1300 1002) was ordered from picodent. Glycerol (cat: 65516-500ml), methanol (cat: 32213-2.5L), protocatechuate 3,4-dioxygenase pseudomonas (PCD) (cat: P8279), 3,4- dihydroxybenzoic acid (PCA) (cat: 37580-25G-F) and (+−)-6-hydroxy-2,5,7,8- tetra- methylchromane-2-carboxylic acid (Trolox) (cat: 238813-5 G) were ordered from Sigma. Potassium chloride (cat: 6781.1) was ordered from Carl Roth. Paraformaldehyde (cat: 15710) were obtained from Electron Microscopy Sciences. 90 nm diameter Gold Nanoparticles (cat: G-90-100) were ordered from cytodiagnostics.

#### <u>B</u>uffers

For imaging the following buffer was prepared: Buffer C (1× PBS, 500 mM NaCl).

Directly before imaging Buffer C was supplemented with: 1× Trolox, 1× PCA and 1× PCD (see paragraph below for details). 100x Trolox: 100 mg Trolox, 430 μl 100 % Methanol, 345 μl 1M NaOH in 3.2 ml H_2_O. 40× PCA: 154 mg PCA, 10 ml water and NaOH were mixed, and pH was adjusted 9.0. 100× PCD: 9.3 mg PCD, 13.3 ml of buffer (100 mM Tris-HCl pH 8, 50 mM KCl, 1 mM EDTA, 50 % Glycerol). All three were frozen and stored at -20 °C.

#### Sample preparation

*Drosophila* hemi-thoraces were isolated and stained as described above with phalloidin Alexa488 (1:2000) and the two nanobodies coupled to either P1, P3 or PS3 oligos (about 50 nM) overnight. Before embedding the samples into the chamber, they were washed two times with PBS + 1% Triton. Hemi-thoraces were embedded as described above. Before assembling the chamber, the cover slip was treated with 90 nm diameter gold nanoparticles (cat: G-90-100, Cytodiagnostics, 1:10 dilution into methanol). After assembling, the chamber was filled with imaging buffer containing the complementary P1, P3 or PS3 imaging oligos (see below for imaging conditions) and sealed with Picodent glue.

#### Super-resolution microscope

Fluorescence imaging was carried out on an inverted microscope (Nikon Instruments, Eclipse Ti2) with the Perfect Focus System, applying an objective-type TIRF configuration with an oil-immersion objective (Nikon Instruments, Apo SR TIRF 100x, NA 1.49, Oil). A 561 nm and 640 nm (MPB Communications Inc., 2 W, DPSS-system) laser were used for excitation. The laser beam was passed through clean-up filters (Chroma Technology, ZET561/10, ZET642/20x) and coupled into the microscope objective using a beam splitter (Chroma Technology, ZT561rdc, ZT647rdc). Fluorescence light was spectrally filtered with an emission filter (Chroma Technology, ET600/50m and ET575lp, ET705/72m and ET665lp) and imaged on a sCMOS camera (Andor, Zyla 4.2 Plus) without further magnification, resulting in an effective pixel size of 130 nm (after 2×2 binning).

#### Imaging conditions

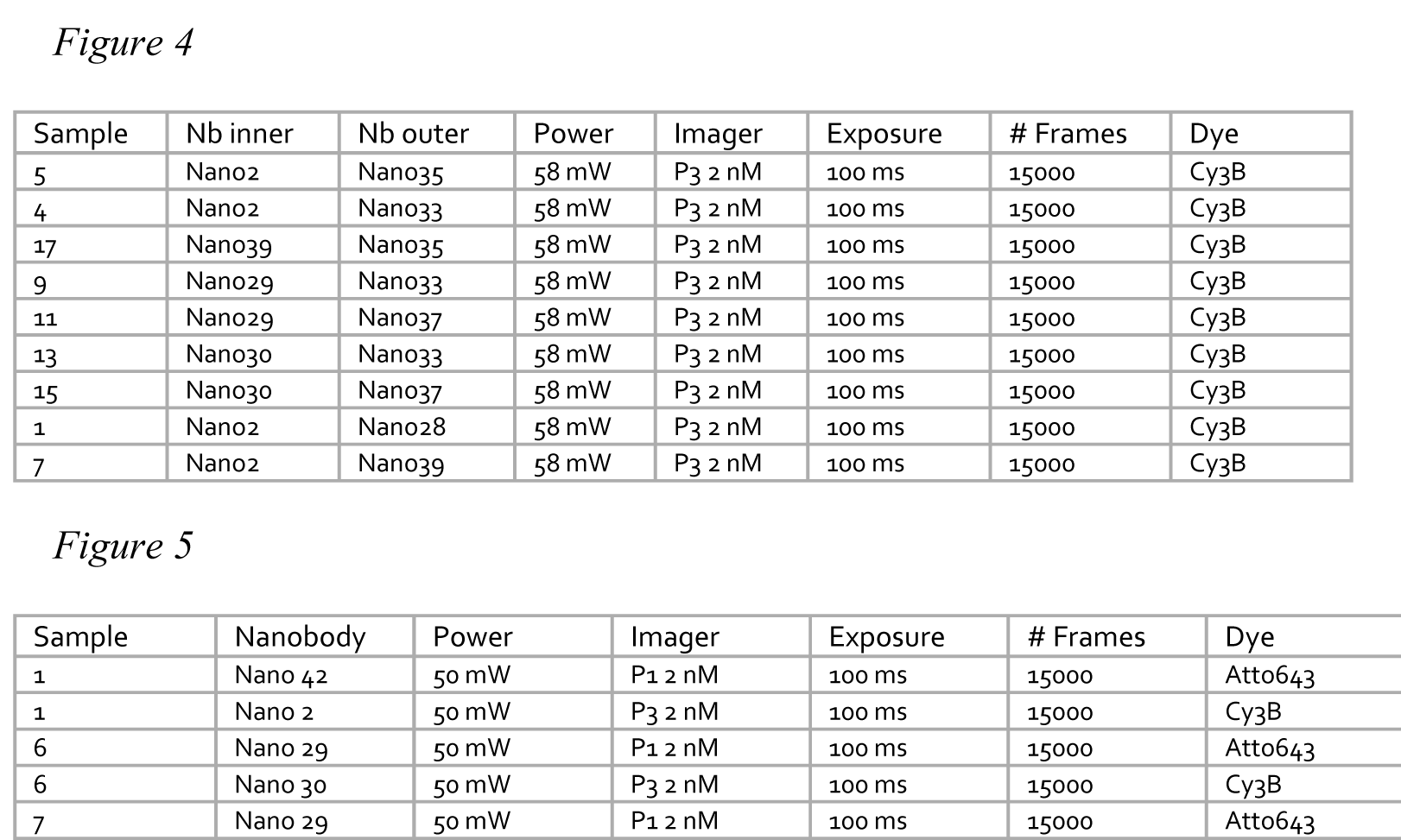
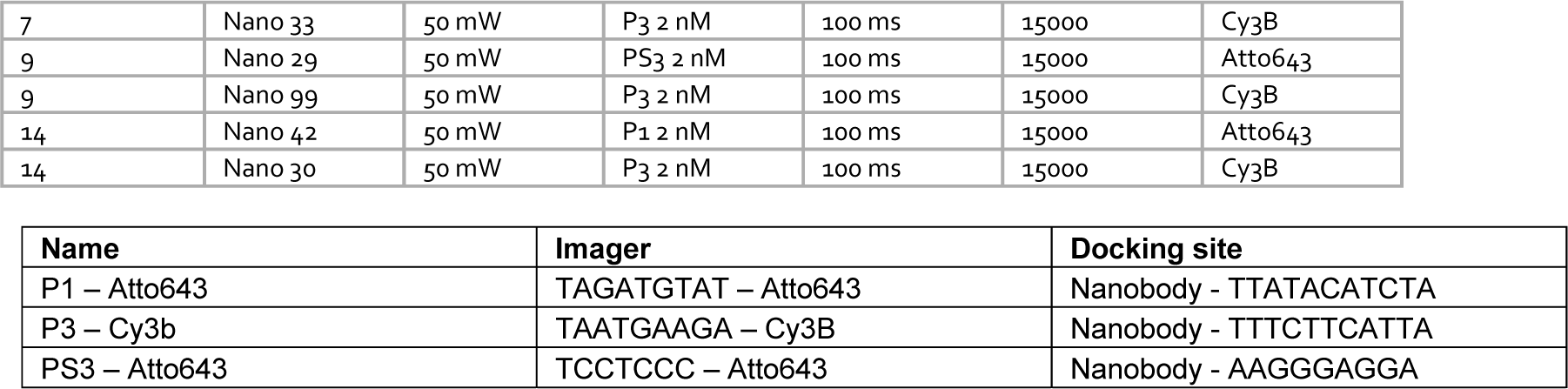

#### Super-resolved image reconstruction

The data acquired during imaging was post-processed using the Picasso (Schnitzbauer et al., 2017) pipeline. First, the localisations were detected by a threshold-based detection and fitted with a least-square fit. Next, the data was drift corrected using a redundant cross-correlation and a fiducial marker-based drift correction. Then, a super-resolved image was rendered using Picasso render. From the images, the myofibrils for further analysis were selected interactively using the rectangular pick tool. All further analysis was done with customized jupyter notebooks.

#### Extraction of band positions from DNA-PAINT data

Extraction of band positions from DNA-PAINT data was achieved the following way: first, individual myofibrils were manually selected using the rectangular selection tool from Picasso (Schnitzbauer et al., 2017) and saved in individual files.

Second, the remaining of the analysis was automated in custom codes written in Python. To limit localisation events arising from multiple emitters that create artefacts (Lelek et al. 2021), localisations were filtered based on the standard deviation of their Gaussian fits. Localisations kept were within a disc in the standard deviation space (sx, sy), centered on the maximum of the distribution and of radius 0.2 pixel.

Third, individual Z-discs were automatically detected. This did not require super-resolved data and the process was the result of multiple steps: (a) the algorithm rotated selections and their localisations to orient the selection horizontally. (b) Localisations were projected along the main axis of the selection and their density was reported in a histogram, which bins size was the same as the pixel size of the camera. The histogram can be seen as a low resolution intensity profile along the myofibril. (c) The algorithm found peaks in the resulting histogram (with find_peaks from the scipy library) corresponding to positions of Z-discs. (d) Once peaks were detected, the algorithm selected peaks that were relevant to the analysis, using the fact that the distance between Z-discs is the size of a sarcomere.

Fourth, with the knowledge of Z-disc positions, the algorithm then focused on windows centered on Z-discs to extract the positions of bands: (a) Similar to step 3, the algorithm rotated the selection and stored localisations in a histogram, which bin size is adjusted for best results (typical bin size was 13 nm). (b) Because DNA-PAINT data accumulate the localisations, the histogram of localisations can display fluctuations that make automated extraction of band positions difficult. Therefore, to locate the rough position of a given band, the data were first convolved with a Gaussian function of standard deviation 25 nm that smoothens fluctuations. (c) The resulting histogram was then analysed with a peak finding algorithm to locate rough band positions. (d) Finally, to precisely locate band positions, the algorithm fitted a Gaussian function on the non-convolved data, in a window centered on each of the positions detected at the previous step. To ensure that the analysis was properly achieved, the results were visually checked.

#### Averaged epitope positions using bootstrapping

To obtain an uncertainty estimate of the averaged position of epitopes, we used the bootstrapping method https://doi.org/10.1007/978-1-4899-4541-9. In brief, each dataset of an epitope is used to create 1000 bootstrap replicates. We generated a replicate by drawing individual values in a given dataset with replacement (*i.e.* each value can be drawn multiple times). The size of one replicate is the same as the one of the initial dataset. From each of these replicates we computed the mean, and therefore obtained 1000 means from 1000 replicates. These 1000 means constitute the bootstrap data presented in Figure 6, each epitope having its own bootstrap data. Finally, 95% confidence intervals were obtained by extracting the 2.5% and 97.5% quartiles from these bootstrap data.

## Acknowledgements

We thank Sandra B Lemke and Aynur Kaya-Çopur for their help in the initial DNA-PAINT pilot experiments. We would like to thank Stefan Raunser and Mathias Gautel and all their group members as well as the Schnorrer and Görlich groups for their stimulating discussions within the StuDySARCOMERE ERC synergy grant. We are indebted to the IBDM imaging facility for help with image acquisition and maintenance of the microscopes.

## Supplementary figure legends

**Figure 1 – figure supplement 1.**
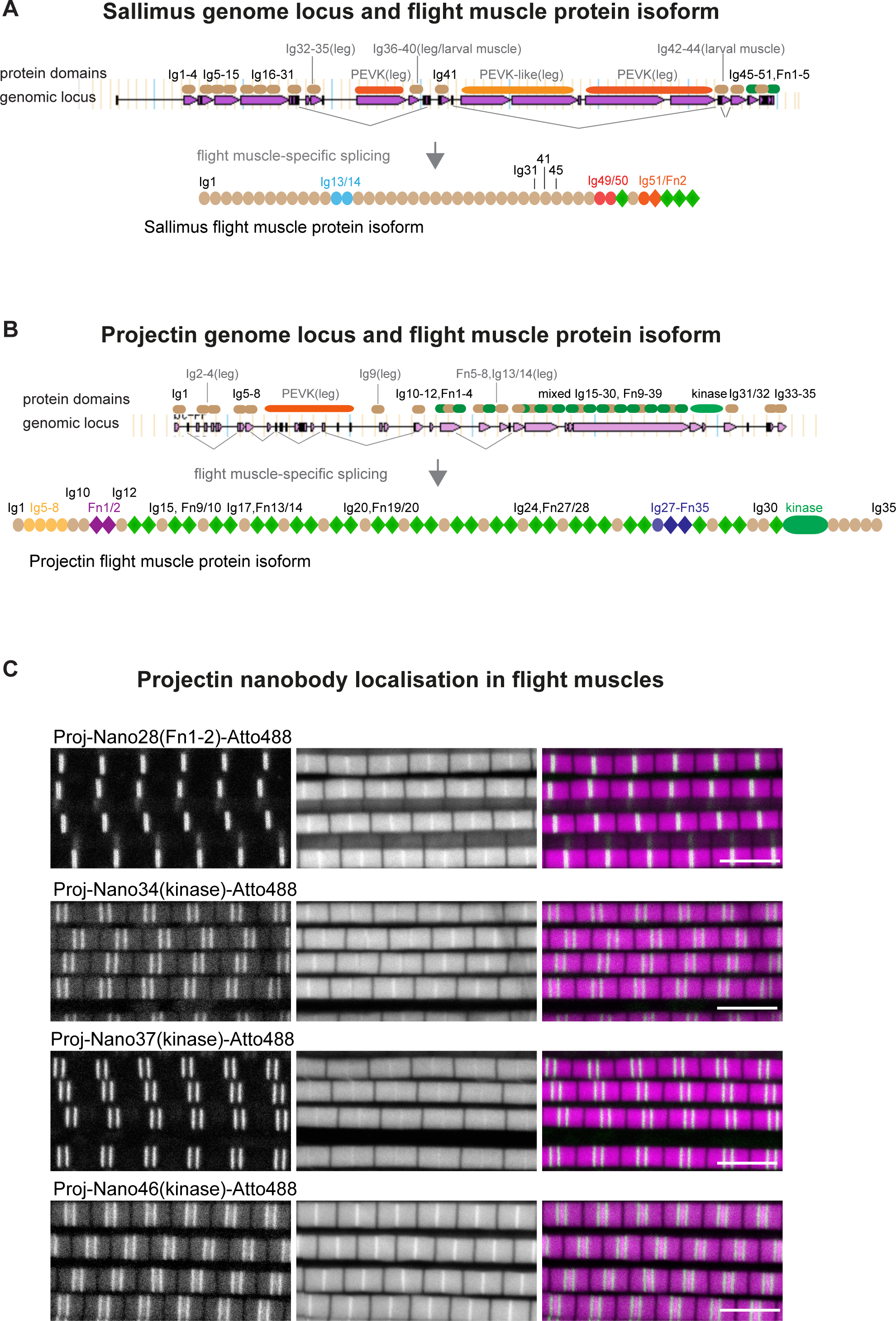
(**A, B**) Genomic organisation of *sallimus* (A) and *projectin/bent* (B) genomic loci taken from Flybase. Pink rectangles indicate exons, lines indicate introns. Above the genomic locus is the rough position of respective protein domains indicated. The domains named in grey are largely specific to leg or larval muscle and not expressed in flight muscles. This is also indicated by the prominent flight muscle-specific splice junctions indicated in grey below the genomic loci. Note that the spring-like PEVK domains in *sallimus* or *projectin* are spliced out in flight muscles. This results in the respective Sallimus and Projection domain structure of the flight muscle isoforms. (**C**) Single confocal sections of flight muscle sarcomeres stained for actin with phalloidin (magenta) and the indicated Proj nanobodies directly coupled to Atto488 (green). The Z-disc is revealed by the prominent actin signal. Scale bars 5 µm.

**Figure 2 – figure supplement 1.**
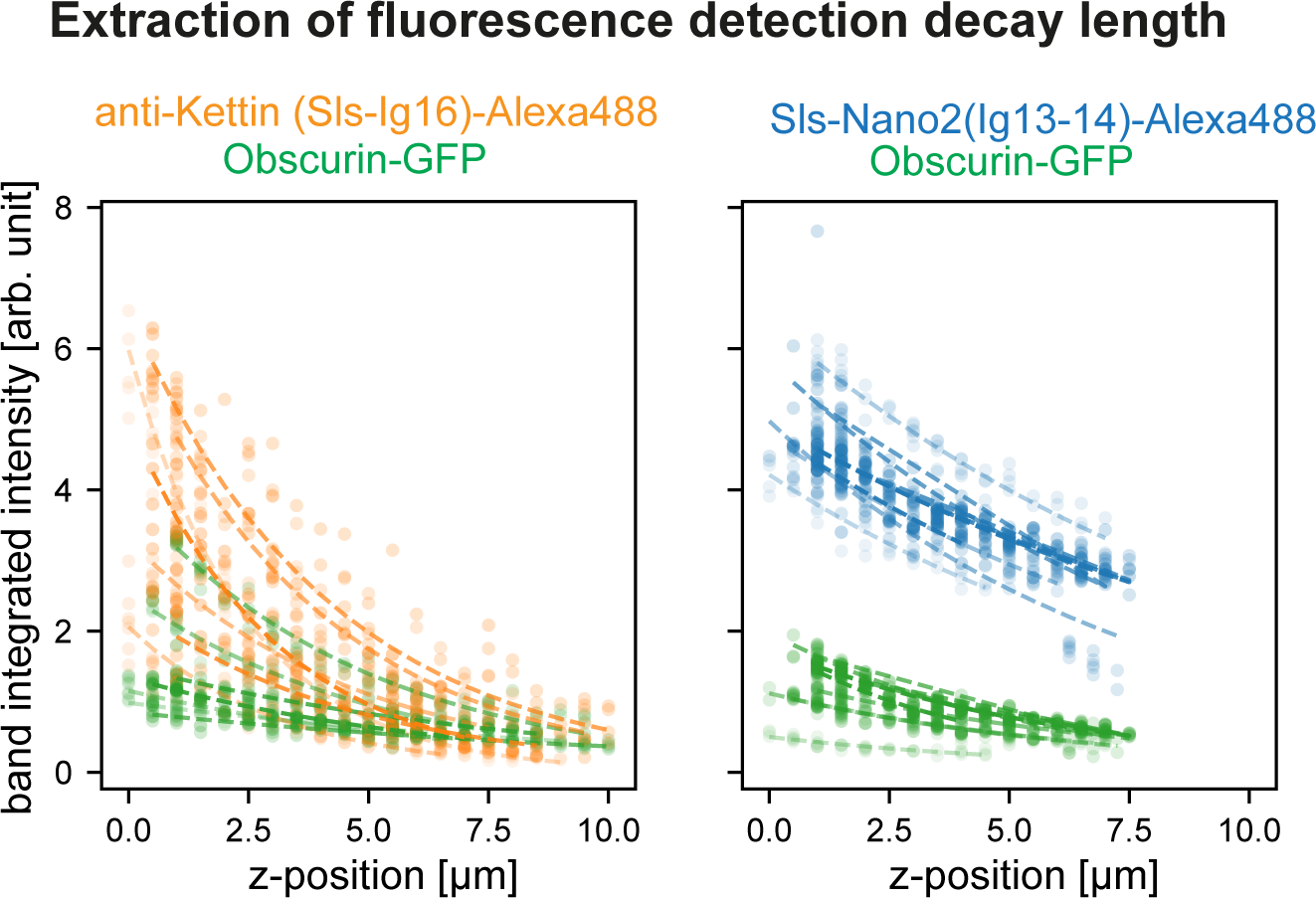
Scatter plots of band mean intensity in a given flight muscle myofibril versus imaging depth. Dashed lines are the exponential decay fits for each hemi-thorax (individual decay lengths obtained from the fits are reported in Figure 2C). GFP in green, anti-Kettin in orange and Sls- Nano2 in blue.

**Figure 2 – figure supplement 2.**
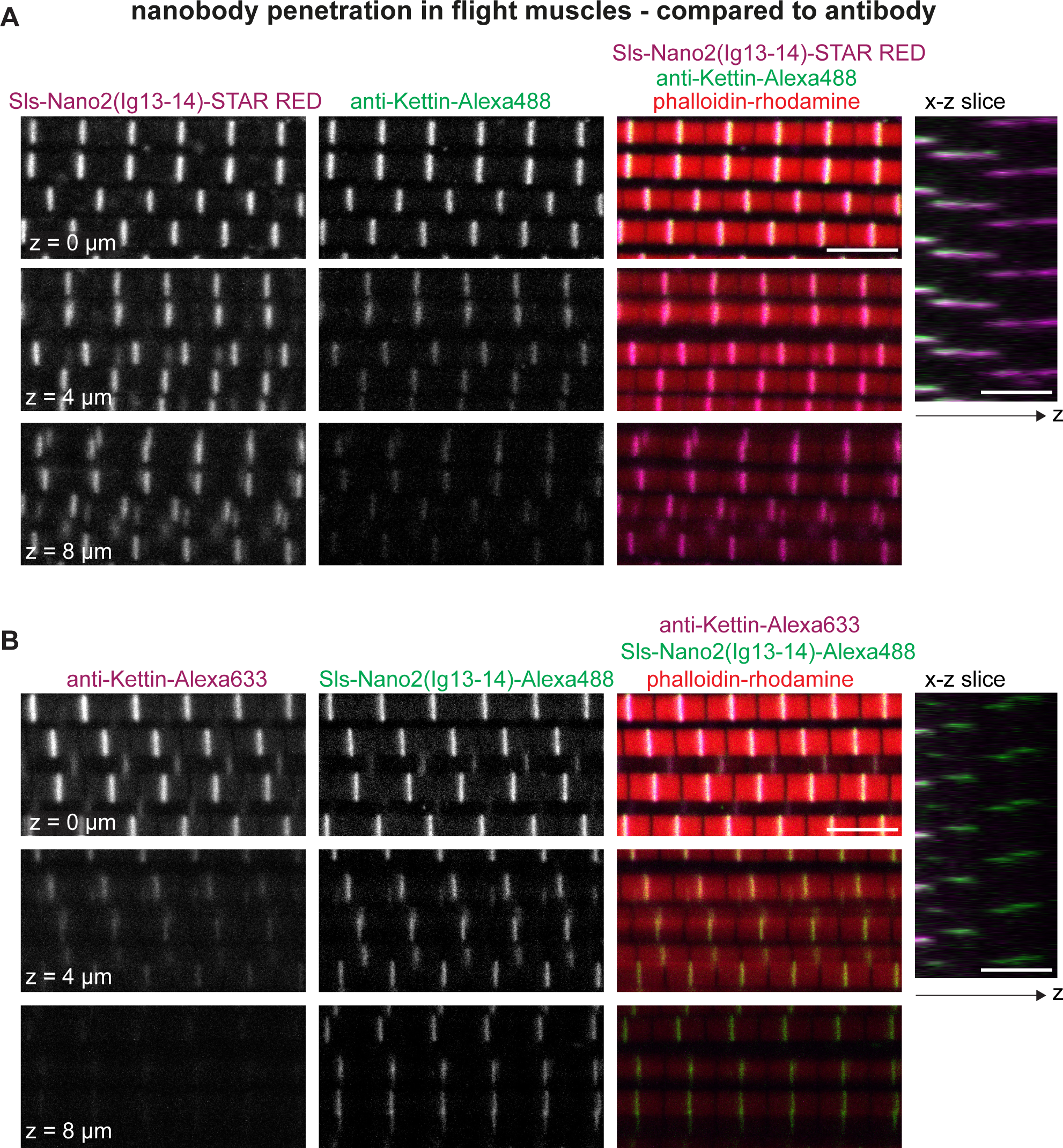
(**A, B**) Flight muscles of adult hemi-thoraces stained with phalloidin (red) and either Sls- Nano2-STAR RED 9 (magenta) together with anti-Kettin antibody and secondary antibody with Alexa488 (green) (A) or Sls-Nano2-Alexa488 (green) and anti-Kettin antibody with Alexa633 (magenta) (B). Three different z-planes and x-z slice are shown. Note that in both examples the nanobody penetrated the entire z-stack, whereas the antibody signal decays quickly in z-direction. Scale bars 5 µm.

**Figure 3 – figure supplement 1.**
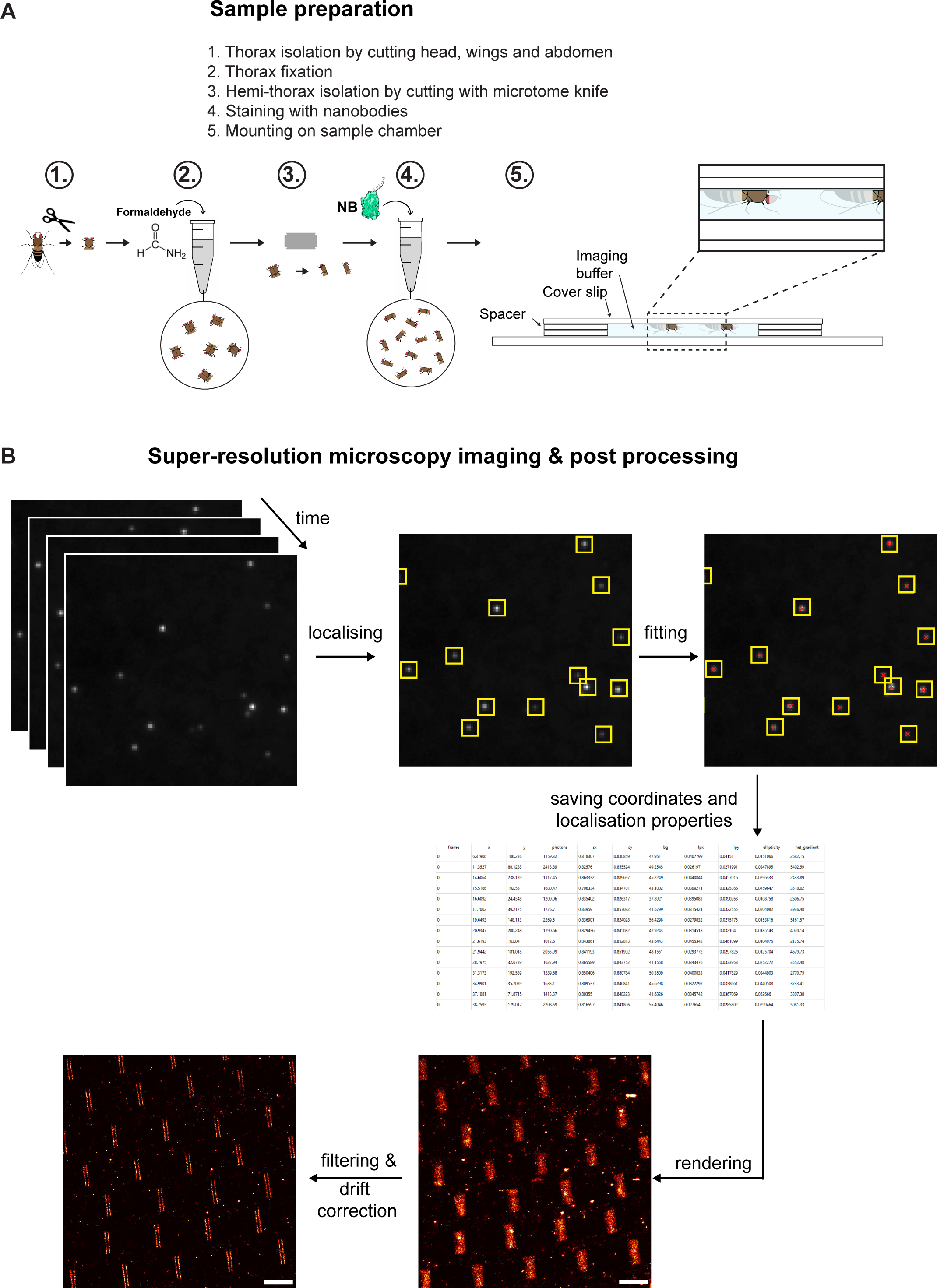
(**A**) Schemes illustrating the different steps of sample preparation. Thorax isolation, followed by PFA fixation and hemithorax preparation. Fixed hemi-thoraces are stained with nanobodies and are mounted in a sample chamber in imager solution positioning the flight muscles close to the coverslip to enable TIRF imaging. (**B**) DNA-PAINT post-processing workflow. The single-molecule ‘blinking’ events are identified using a threshold detection and the maximum is determined via a Gaussian fit. The coordinates in time and space and other localisation properties are saved in a hdf5 file. The hdf5 is then drift corrected and rendered using Picasso render.

**Figure 3 – figure supplement 2.**
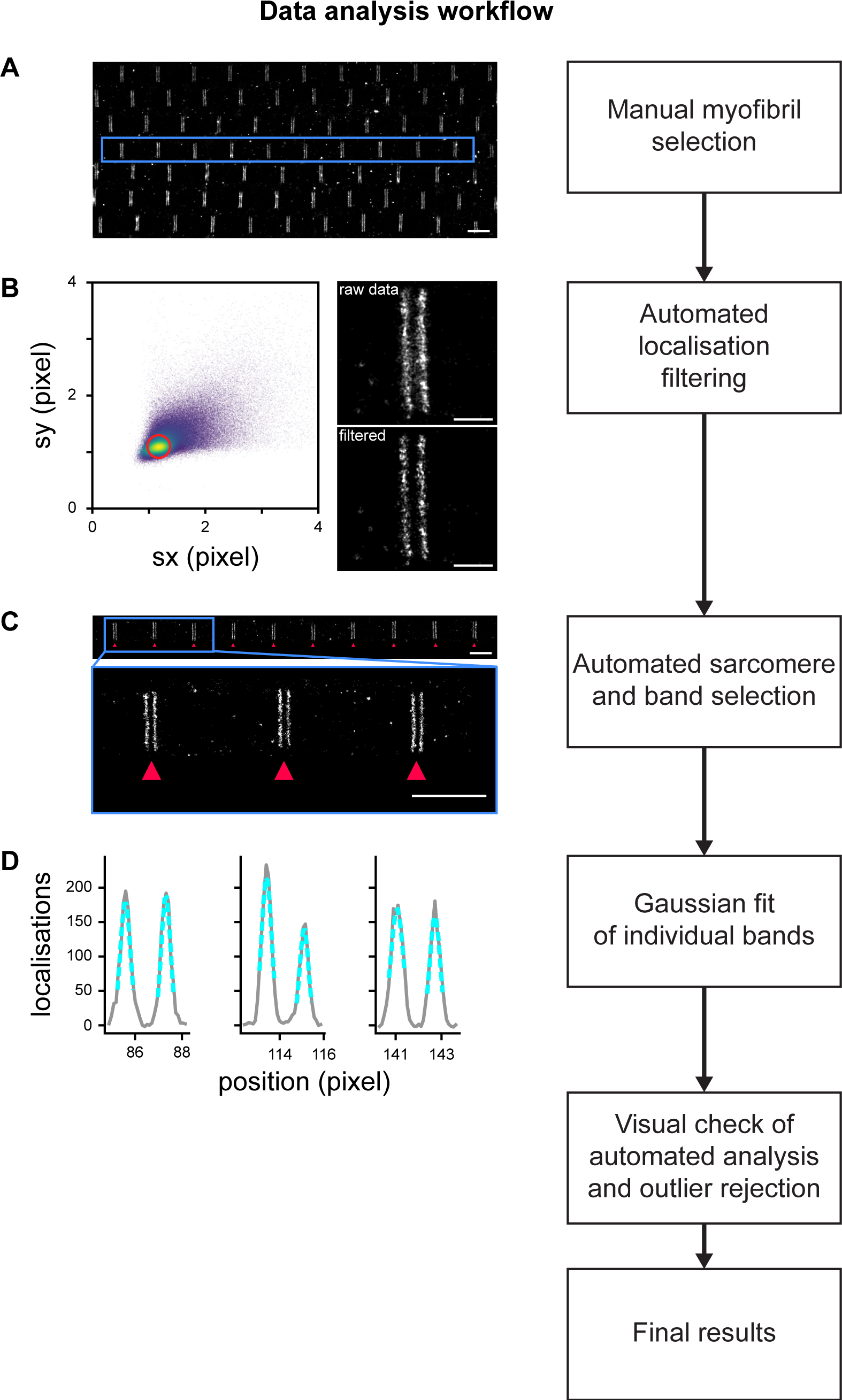
Data analysis workflow (**A**) Individual myofibrils are manually selected using the rectangular selection tool from Picasso (Schnitzbauer et al., 2017). Scale bar 2 µm. (**B**) To limit localisation events arising from multiple emitters that create artefacts (Lelek et al., 2021), localisations are filtered based on the standard deviation of their Gaussian fits. Localisations kept are within a disc in the standard deviation space (sx, sy), centered on the maximum of the distribution and of radius 0.2 pixel. Scale bars 0.5 µm. (**C**) Individual Z-discs are automatically detected (see Methods for details). Scale bars 2 µm. (**D**) Individual bands generated by the accumulation of protein epitopes are detected automatically and their center position is obtained using a Gaussian fit. After a visual check of the automated detection result, the results are compiled for further analysis.

**Figure 5 – figure supplement 1.**
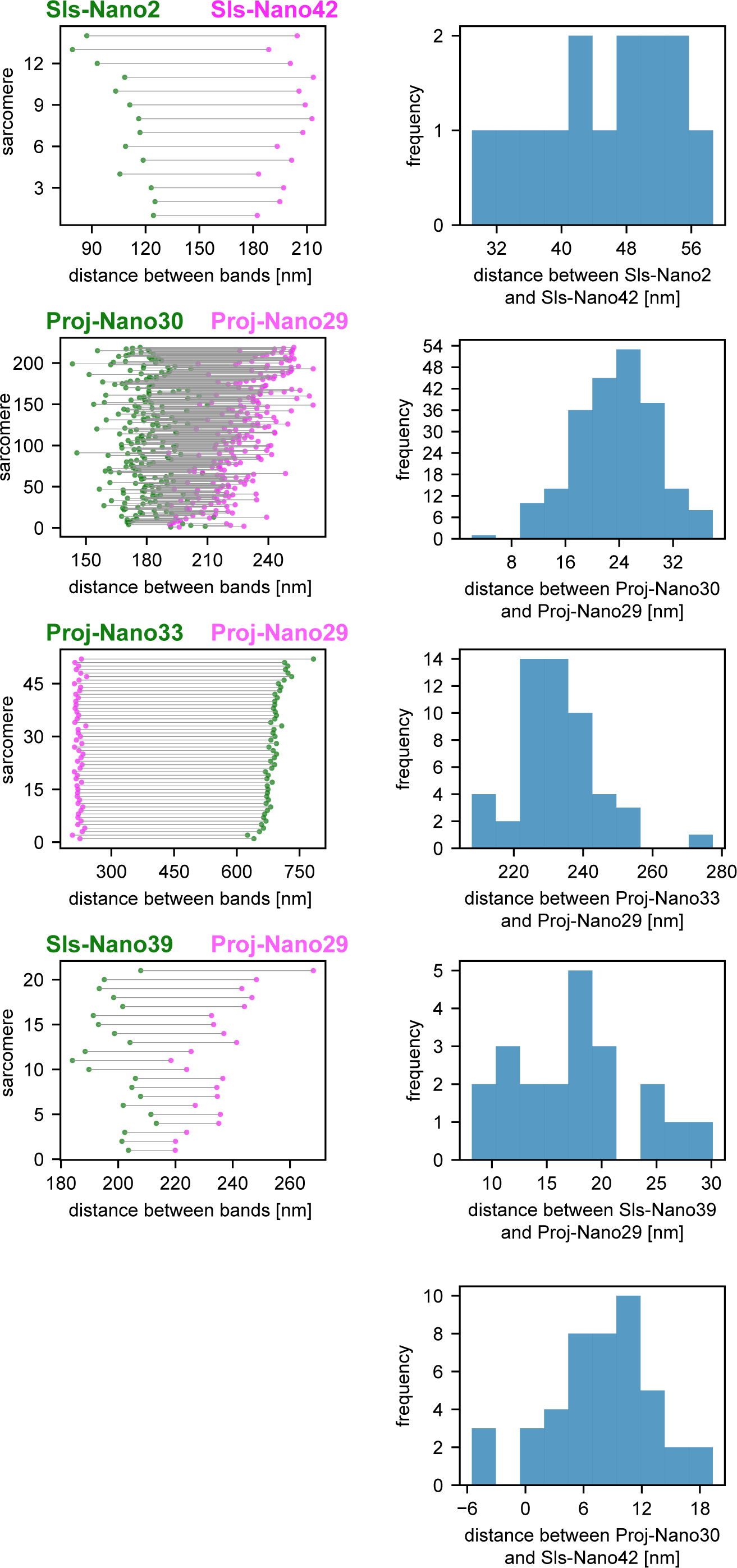
Dual-colour DNA-PAINT sarcomere quantifications Distance quantifications of the dual-colour DNA-PAINT data shown in Figure 5. Plots on the left show the epitope distances from the Z-discs in each individual sarcomere analysed with green and magenta colours indicating the respective nanobodies used. Plots on the right display histograms plotting the distances in between the individual two nanobody epitopes in each sarcomere. These are half the values from the plots on the left, as the organisations are symmetric around the Z-discs.

**Figure 6 – figure supplement 1.**
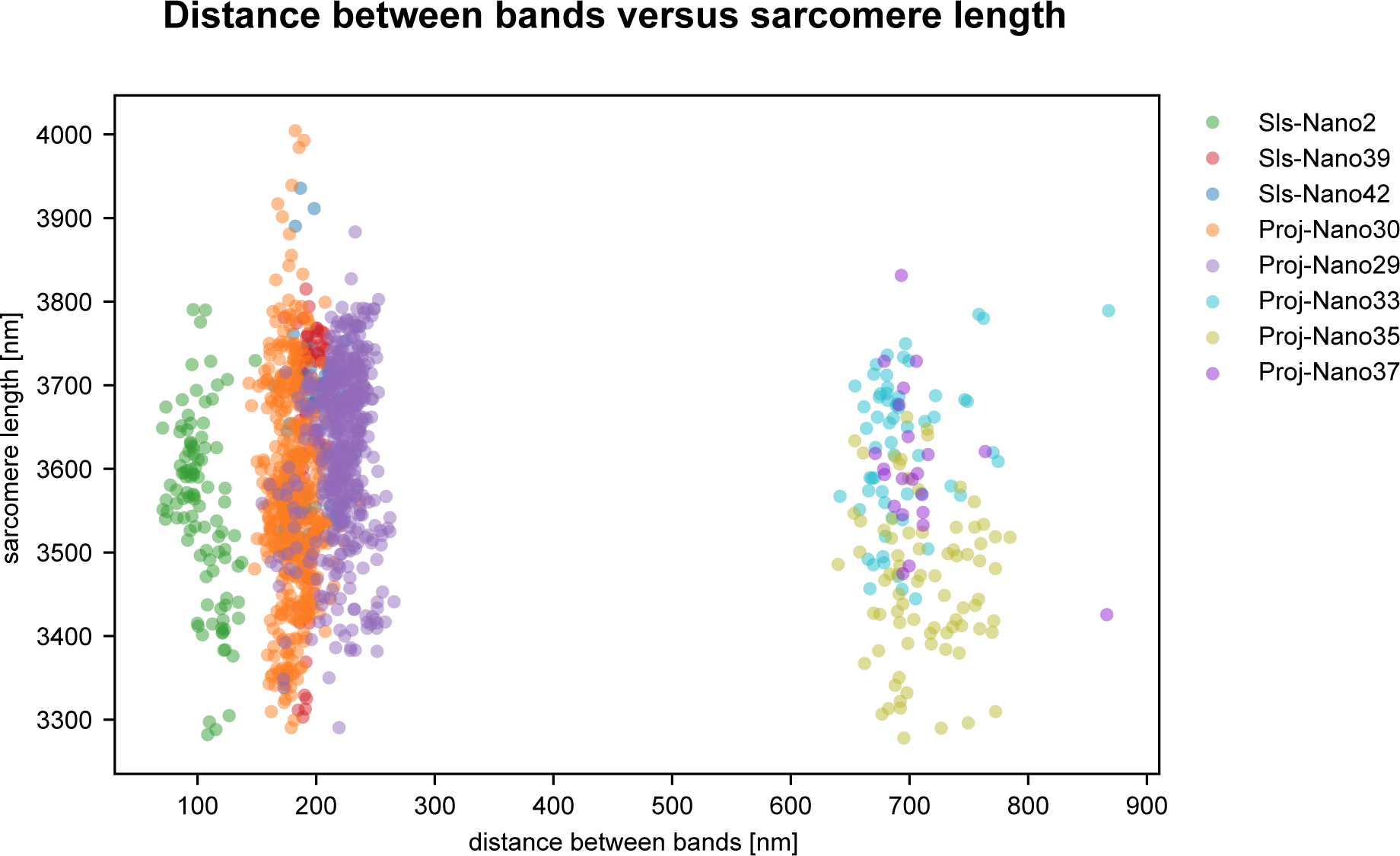
Distance between bands versus sarcomere length Scatter plot of distance between bands of nanobodies used in this study versus the individual sarcomere length. The sarcomere length reported for nanobodies centered on a given Z-disc is obtained by measuring half the distance between Z-discs located directly on the left and on the right. Note that the distances between bands do not correlate with the slight variations in sarcomere length.

